# Mass Spectrometric Determination of Site-Specific O-Acetylation in Rhamnogalacturonan-I Oligomers

**DOI:** 10.64898/2025.12.12.694011

**Authors:** Liyanage Devthilini Fernando, Xu Yang, Stephanie Archer-Hartmann, Christian Heiss, Parastoo Azadi

## Abstract

O-Acetylation, a common modification in rhamnogalacturonan I (RG-I), is critical for various biological processes, including plant growth, stress responses, and pathogen defense. Precise determination of the degree and specific positions of acetylation is therefore essential. To date, nuclear magnetic resonance (NMR) and tandem mass spectrometry have been employed to identify acetyl positions in pectin oligosaccharides. Although NMR is effective, it requires pure, high-concentration samples. Tandem mass spectrometry (MS), which uses lower sample amounts, faces challenges due to acetyl migration between monosaccharide positions. The multiple steps in pectin sample analysis can further promote O-acetyl migration, especially near free hydroxyl groups. Moreover, during tandem MS, acetyl groups may detach, complicating accurate tracking. This study presents an approach to lock O-acetyl groups by introducing trideuteroacetyl and propionyl substituents onto free hydroxyls of RG-I or partially acetylated RG-I. By combining matrix-assisted laser desorption/ionization-time of flight (MALDI-TOF) MS and electrospray ionization (ESI) MS with MS/MS or tandem mass spectrometry (MS^n^), we devised a way to determine the monosaccharide sequence in the oligomer and precise positions of acetyl groups in partially acetylated RG-I. This method enables the study of the regiospecificity of recombinant pectin O-acetyltransferases and can be applied to other oligosaccharides to determine acyl positions.

**GRAPHICAL ABSTRACT:** 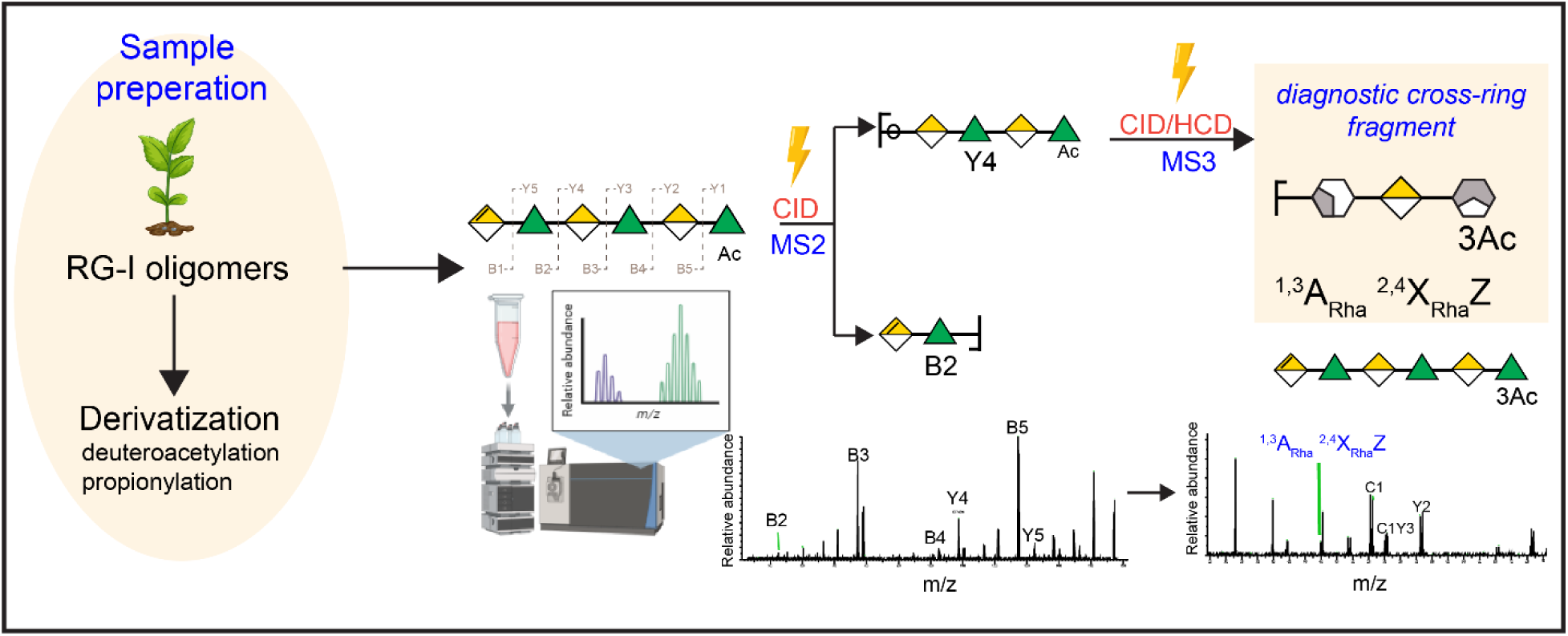

## INTRODUCTION

Rhamnogalacturonan I (RG-I) is a structurally complex pectic polysaccharide that is present in the cell walls of all vascular plants.^1–3^ Due to its structural complexity, little is known about RG-I structural diversity in different cells, tissues, and species, or about how it interacts with cellulose and other polysaccharides to regulate the properties and functions of the wall. O-Acetylation (OAc) is a common non-carbohydrate modification observed in pectin RG-I and plays crucial roles in various biological processes such as growth of plants,^4, 5^ abiotic stress responses,^6^ and defense pathways against pathogens.^7^ To study these processes on a molecular level, it is essential to accurately determine both the overall degree of O-acetylation and the specific positions of acetyl groups.

To date, both nuclear magnetic resonance (NMR) and tandem mass spectroscopy (MS^n^) have been employed to identify the acetylated positions in the pectin oligosaccharides.^8–17^ While NMR is a powerful tool, it requires pure samples and high concentrations. In contrast, tandem mass spectroscopy (MS^n^) can precisely locate acetylation and methylation in oligogalacturonides derived from enzymatic hydrolysis of pectin using very low sample amounts (5 µg).^15–17^ Nevertheless, this task is hampered due to the tendency of acetyl groups to migrate between different positions on a given monosaccharide residue,^8, 18^ multiple isolation and analysis steps involved in studying pectin samples provide ample opportunities for O-acetyl migration, particularly in the presence of nearby free hydroxyl groups. Further, when performing tandem MS, acetyl groups can be lost, preventing the detection of a diagnostic fragment and hindering accurate position tracking.^19–21^

In previous studies, several attempts have been made to prevent migration of acetyls and to determine their location in the oligosaccharides by derivatizing the sample using propionylation^22^,trideuteroacetylation,^23^ and neutral permethylation.^24^ Later, Sharp et al. demonstrated that collision-induced dissociation (CID) could effectively determine the position of *O*-acetyl groups in partially methylated and partially acetylated oligosaccharides (chondroitin sulphate and heparan sulphate).^25, 26^ Derivatization not only fixes the acetyl groups but also enhances the ionization efficiency while lowering the detection limit, particularly in higher order MS^n^ (n>2) experiments.^27^ However studies focusing on the precise determination of O-acetylation sites in the pectin oligosaccharides especially those derived from RG-I remain scarce.

Therefore, in this study, we have successfully developed a method to lock the natively present O-acetyl groups in place by introducing trideuteroacetyl and propionyl substituents onto free hydroxyls of partially acetylated RG-I oligosaccharides. This work is built upon previous derivatization methods, with the aim to overcome the shortcoming of these promising but not yet fully mature methodologies and render them more broadly applicable. Using a combination of matrix-assisted laser desorption/ionization time-of-flight mass spectroscopy (MALDI-TOF MS), electrospray ionization mass-spectroscopy (ESI MS) with MS/MS or MS^n^ tandem mass capabilities, and liquid chromatography (LC) separation with MS^n^ schemes allows us to determine the structure of the oligomers and the precise position of acetyl group in partially acetylated RG-I. This method can be used to examine the regiospecificity of recombinant pectin O-acetyltransferases and can be extended to other oligosaccharides to determine the acyl positions.

## MATERIALS AND METHODOLOGY

Acetic anhydride-d_6_ (99 atom %D) was obtained from Sigma-Aldrich (St. Louis, MO, USA). Pyridine was purchased from J.T Baker (Phillipburg, NJ, USA) propionic anhydride. Dichloromethane (DCM) used for extraction of derivatized sample was purchased from Sigma-Aldrich. The 2,5-dihydroxybenzoic acid (DHB) matrix was purchased from Sigma-Aldrich, MO, USA. The ESI-MS solvent acetonitrile (ACN) was purchased from Honeywell (Tokyo, Japan). RG-I (*Arabidopsis mucilage*) partially acetylated oligosaccharide and digested RG-I celery samples were obtained from the Dr. Breeanna Urbanowicz laboratory at the Complex Carbohydrate Research Center, University of Georgia.

### Sample preparation

#### Enzymatically acylated RG-I degree of polymerization 6 (DP6)

Pectin lyase digested *Arabidopsis mucilage* RG-I oligomers (average degree of polymerization 6) was enzymatically acetylated using trichome51 birefringence (TBR) enzyme using 4-methulumbelliferyl acetate as the donor in the MES-buffer at 30 ℃ as reported previous.^34^ The acetylated RG-I product was separated using Superdex peptide 300/10 GL column (cytiva) connected to Agilent 1260 infinity II HPLC. The 10 mM ammonium acetate was used as the mobile phase at the 0.5 ml/min rate. The RG-I containing fractions was lyophilized and further analyzed with NMR and mass spectroscopy.

#### Native celery RG-I

The celery RG-I oligomers were prepared using the BT4175 RG-I lyase enzyme as reported previously.^38^ It cleaves the backbone and generates an unsaturated double bond between C4 and the C5 of the non-reducing GalA residues in the RG-I backbone.^38^ The digested celery RG-I oligomers were directly used for derivatization and analysis.

### Derivatization

The peracetylation procedure was modified from Xu et al 2019^31^. RG-I pyridine and acetic anhydride-d6 (1:1) was reacted at 60 °C for 24 h. The propionylation was carried out in the same way with pyridine and propionic anhydride (1:1) at 60 °C for 24 hrs. The reaction product was extracted with DCM and washed five times with water. The DCM layer was dried under dry air. The samples were dissolved in acetonitrile prior to the mass spectroscopy analysis.

### MALDI-TOF

Samples were dissolved in acetonitrile or methanol water 1:1 mixture (∼final concentration 20mg/mL) for MALDI-TOF analysis. 1 µL of sample was spotted on a ground steel MALDI plate with 1 µL of 2,5-dihydroxybenzoic acid (DHB) matrix (15 mg/ml; in 70% ACN:0.1% formic acid) and after crystallization, the spot was analyzed on a Brucker rapfleX Tissuetyper MALDI-TOF MS instrument (Bruker Daltonics, Germany) under reflector mode in positive or negative ionization mode. The mass spectra were processed for smoothing and baseline subtraction using FlexAnalysis version 3.4 (Bruker Daltonics, Germany). For MS2 analysis on the MALDI-TOF instrument, the targeted ions were isolated and subjected to collision-induced dissociation (CID) for fragmentation analysis.

### Direct Infusion (DI) ESI MS^n^ analysis of RG-I DP6

Experiments utilizing direct infusion (DI) ESI-MS^n^ on derivatized RG-I oligosaccharides were performed on an Orbitrap Eclipse Tribrid mass spectrometer with both positive and negative polarities as needed.

Derivatized sample solutions were diluted to 20-50 µg/ml solution with acetonitrile. Direct infusion was conducted at a flow rate of 3 µl/min. The EASY-Max NG heated ESI (H-ESI) source used in experiments was supplied by nitrogen generators. A HESI-II probe with low-flow needle insert was used for generating highly sensitive MS^n^ spectra. The ion transfer tube was maintained at 325 ℃, the vaporizer was held at 52 ℃. The spray voltage and aux gas, sheath has and sweep gas flow rates were adjusted for optimal signal intensity and stability. For precursor isolation quadrupole was used, while radio frequency (RF) lens is the most critical factor directing the enhancement of ions at different mass ranges. Most precursor ions of different levels of MS^n^ were able to be fragmented by 30-45% (normalized collision energy) CID. However, some of the derivatized RG-I fragments were resistant to 100 % CID fragmentations. Further higher-energy collision dissociation (HCD) was used for MS^3^ fragmentation to obtain cross-ring fragmentation. A total of 100 scans were averaged for each experiment to improve the data quality.

### LC-ESI MS^n^ analysis of Celery RG-I

A derivatized RG-I sample from celery was subjected to LC-MS^n^ analysis on Ultimate 3000 RSLCnano system coupled with Orbitrap Fusion Tribrid mass spectrometer (ThermoScientific). A Commercial Acclaim Pepmap100 C18 0.75 × 150 mm column operated at 55 °C was used for sample separation. Buffer A was 0.1% formic acid; Buffer B was 0.1% formic acid in 80% acetonitrile. The elution gradient was 10-95% B in 40 min after the column was equilibrated by 10% B for 10 min at the beginning of each injection. Ten microliters of sample were injected. The targeted MS^3^ spectra was recorded in parallel with a data-dependent acquisition MS^2^ program for precursor screening. A 3 Da isolation window was applied for 45% CID MS^2^ and a 4 Da isolation window for 35% HCD MS^3^. Quadrupole isolation was used, and the RF lens was set at 60%. The target MS^n^ list was prepared based on direct infusion observations and theoretical masses calculated using GlycoWorkbench 2.1 and ChemDraw 23.0.

### QExactive analysis of Celery RG-I

The native RG-I celery sample was analyzed using Vanquish Ultra Performance LC (UHPLC) (ThermoFisher Scientific) connected to a Thermo QExactive HF orbitrap mass spectrometer (ThermoFisher Scientific). The separation was carried out on a commercial C18 column (Zorbax Eclipse XDB-504C18; 2.1x150 mm; 1.8 μm) with a linear gradient from 1 % to 99 % acetonitrile in water containing 0.1 % formic acid at a flow rate of 0.3 ml/min 22 minutes. The column was cleaned for 2 minutes with 99 % acetonitrile followed by 4 minutes of reconditioning with 1 % acetonitrile. A data-dependent program was used for acquisition in the positive ion mode, where the precursor ion scan (MS1) was acquired at 120k resolution from 200-2000 followed by top-down fragmentation (stepped HCD; MS2) of high-to-low intensity m/zs at 30k resolution.

### NMR analysis of RG-I

Approximately ∼0.5 -1 mg of the RG-I DP6 oligomer and RG-I celery was dissolved in 200 µL of D_2_O (99.9 % D) and lyophilized overnight. Then the sample was dissolved in 40 µL of D_2_O (99.96 % D). The dissolved sample was transferred to a 1.7-mm NMR tube for NMR analysis. NMR data were acquired at 25 ℃ on a Bruker Avance NEO spectrometer (^1^H, 800 MHz) equipped with a cryoprobe using standard pulse sequences. The acquisition parameters are tabulated below in **Supplementary Table 1**. Chemical shifts were referenced to 3-(trimethylsilyl)-propanesulfonic acid sodium salt (DSS-d_6_) peak (δ_H_ = 0.0 ppm, δ_C_ = 0.0 ppm), for RG-I DP6 oligomer and for RG-I celery referenced to HOD peak (δ_H_ = 4.78 ppm). The spectra were processed and analyzed with *MestReNova* v14.2.1-27684.

### Data Analysis

The general method on acetylation position determination is summarized in **Supplementary Methods 1**. The ESI-MS^n^ data obtained from the Orbitrap Tribrid mass spectrometer were analyzed using Free style software version 1.8 SP1(Thermo Fisher Scientific). The MALDI-TOF MS/MS spectra were analyzed using Brucker flex ABakysis (version 4.2). The fragmentation patterns were predicted using the Glyco Workbench (version 2.1) for native and deuteroacetylated RG-I and ChemDraw (version 23.0.1.10) was used to predict the propionylated RG-I.

## RESULTS AND DISCUSSION

### Derivatization of RG-I

To develop a robust method for determining the acetyl position in the RG-I, the first step was to find the best way to derivatize the samples for the purpose of blocking all free hydroxyls to prevent migration of the acetyl group. Initially, we carried out derivatization using different acylating reagents on neutral (maltohexose) and acidic (RG-I) oligomers to find the optimum conditions to get fully acylated oligomers. We used slightly modified reported Methods **1-3** (**Table 1**,) either to acetylate or propionylate the oligomers. Method **2** gave fully derivatized neutral and RG-I oligomers but Method 1 yielded incomplete derivatization (**Supplementary Figure 1**). Method **3** (perchloric acid) was too harsh even for neutral oligomers and led to sample degradation (**Supplementary Figure 1**). The pyridine catalyst (in Method **2**) typically allows for more selective and milder and neutralizes the byproducts^28^ and was also used previously to acylate pectin.^29, 30^ Therefore, we used pyridine with the appropriate acid anhydride as the acylating reagent in this study.

**Table 1:** Acylating reagents used to derivatize.

| Method | Acylating reagent | Reaction Conditions | Reference | Outcome |
| --- | --- | --- | --- | --- |
| 1 | Trifluoroacetic anhydride and appropriate acid (propionic acid, deuterioacetic acid) | 24 h at 60 °C | Manzi et al 1990<sup>23</sup> | Partial derivatization |
| 2 | Pyridine and propionic anhydride/acetic anhydride | 24 h at 60 °C | Xu et al 2019<sup>31</sup> | Fully Derivatized |
| 3 | Propionic anhydride/perchloric acid |  | Zhong et al 2020<sup>32</sup> | Too harsh |
| 4 | Methyl trifluoromethanesulfonate in tri-methyl phosphate | 2 h at 50 °C | Prehm et al 1980 | Not successful |

We applied the pyridine trideuteroacetic anhydride/propionic anhydride derivatization condition (Method **2**) to the native unsaturated RG-I oligomer (unsaturated uronic acid residue at the non-reducing end of the oligomer) with degree of polymerization 6 (DP6, molecular weight MW= 966 Da) obtained from *Arabidopsis* mucilage by lyase digestion (without O-acetyl groups). This method gave full acylation of the RG-I oligomer (**Supplementary Figure 2**). The acylation reaction used a 1:1 mixture of pyridine: acetic anhydride-d_6_ or propionic anhydride at no more than 60 ℃ to prevent decomposition of the RG-I molecule. The reaction product was then extracted using dichloromethane, as the acylated RG-I is more hydrophobic than native RG-I. Based on the MALDI-TOF MS spectra, the product of the acylation of RG-I DP6 was 80-90% deuteroacetylated or propionylated. The acyl groups substituted all the 2-OH and 3-OH of galacturonic acid (GalA) and 3-OH and 4-OH of rhamnose (Rha) in the RG-I oligomer.

We also attempted methylation of RG-I oligomers by Method **4,** using methyl trifluoromethanesulfonate in trimethylphosphate as reported in Prehm et al 1980 (**Supplementary Figure 3**). However, this reaction condition was ineffective for the RG-I oligosaccharides, likely because of limited solubility of acidic oligosaccharides in trimethylphosphate. The lower pH and presence of moisture coming from the reagents may also have played a role in preventing methylation.

Based on the MALDI-TOF MS and MS/MS of the deuteroacetylated and propionylated samples the respective singly charged [M+Na] ^+^ parent ion (*m*/*z* =1574 and 1717) yielded, upon collision, numerous daughter ions characteristic of these oligosaccharides (**Supplementary Figure 2**). The major ions are result of B and Y cleavages providing information related to the monosaccharide sequence of the RG-I oligosaccharide. The conversion of the RG-I oligosaccharides to their corresponding derivatives improved the ionization efficiency and detection sensitivity by one to over two orders of magnitude compared with the detection of non-derivatized analytes and allowed for detection in lower concentrations (e. g. nanomolar, picomolar levels). When compared to the native RG-I, deuteroacetylation increases the mass by 45 Da per acyl group, and propionylation increases the mass by 54 Da per acyl group. This derivatization was reproducible and consistently yielded derivatized DP6, DP8 and DP10 RG-I (**Supplementary Figure 4**).

### Stability of O-acetyl group in the acylation condition and acetyl position determination using MALDI-TOF-MS

To ascertain that native O-acetyl groups are stable under the deuteroacetylation and propionylation reaction conditions, we exposed enzymatically partially (mono, di) acetylated RG-I DP6 molecules to the acetylating reagent (deuteroacetic anhydride/propionic anhydride + pyridine) and performed MALDI-TOF-MS on peracylated samples (**Figure 1A and 1C**). Both deuteroacetylated and propionylated RG-I showed a mixture of non-acetylated, mono-acetylated and diacetylated RG-I DP6 in the sample.

**Figure 1:**
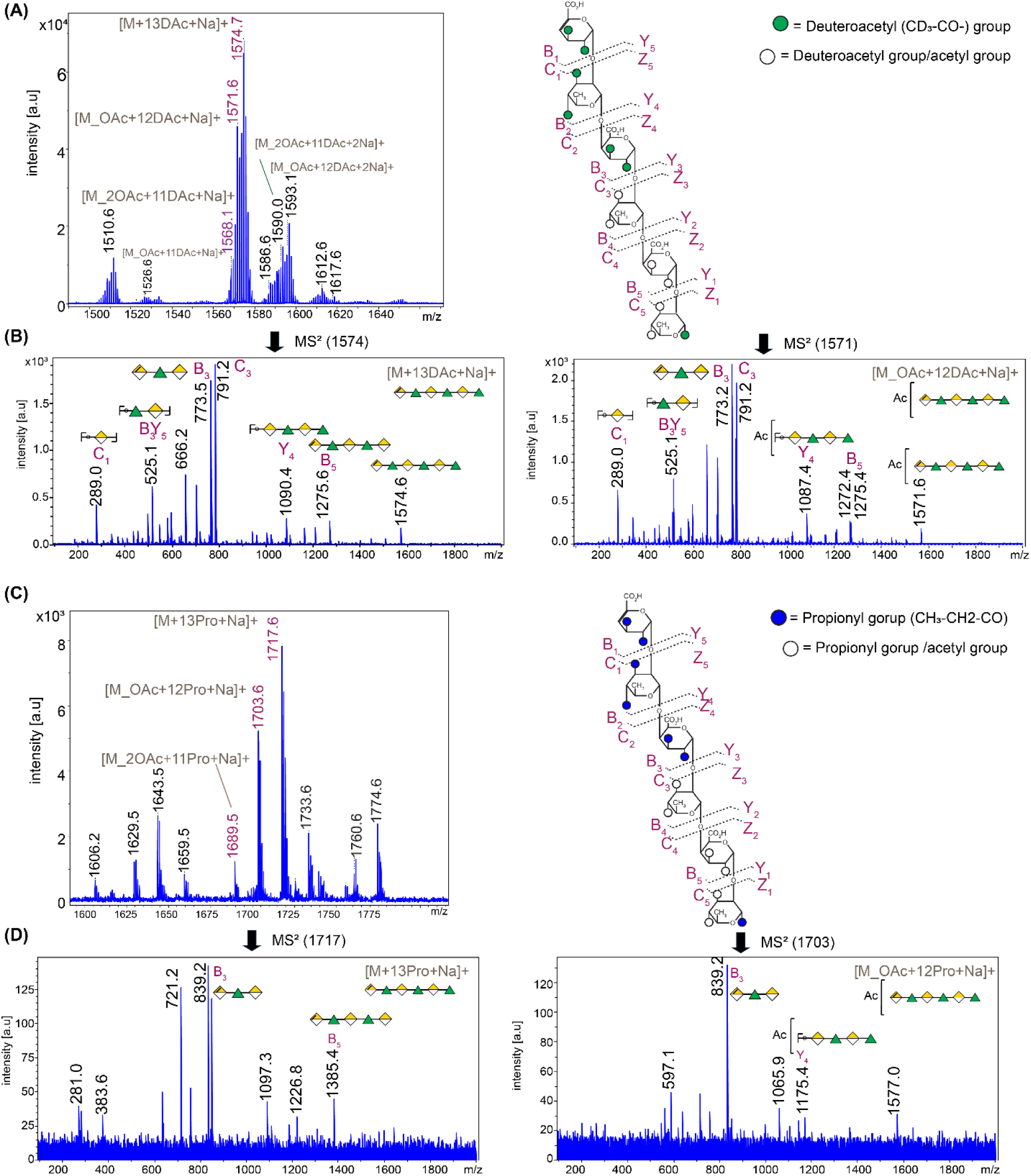
Derivatization of partially acetylated RG-I (RG-I OAc DP6). **(A)** Deuteroacetylated RG-I MS^1^ spectrum (left), RG-I structure, green circles denote deuteroacetyl groups white circles denote either deuteroacetyl or acetyl groups. **(B**) MS2 spectra for *m/z*=1571 [RG-I_OAc DP6+Na]^+^ and *m/z* =1574 precursor ion corresponds to [RG-I DP6+Na]^+^. **(C)** Propionylated RG-I DP6 MS^1^ spectrum (left), RG-I structure, green circles denote deuteroacetyl groups white circle denotes either deuteroacetyl or acetyl groups (right). **(D**) MS2 spectra of *m/z* =1717 [RG-I DP6+Na]^+^ and *m/z*=1703 [RG-I_DP6OAc+Na]^+^. Fragmentation nomenclature shown according to Domon and Costello^33^ for RG-I. Based on the MALDI-TOF spectra the whites circles shows the possible positions of acetyl groups. The non-labelled peaks, the structural composition is not deducted.

Following perdeuteroacetylation, the mono-acetylated RG-I oligomer (*m/z* =1571) gave a mass of 3 Da lower than its non-acetylated counterpart (*m/z*=1574) indicating that the acetate group was not replaced by deuteroacetate (**Figure 1A**). Similarly, in the propionylated RG-I, a 14-Da difference was observed between the non-acetylated and the mono-acetylated RG-I oligomers (**Figure 1C**). Furthermore, we qualitatively assessed the peak ratio of acetylated to non-acetylated RG-I species before and after derivatization using the MALDI-TOF spectra. The ratios remained approximately the same even after the derivatization (60% of non-acetylated RG-I and 40% of mono-acetylated RG-I), confirming that the acetyl groups were stable under the deuteroacetylation or propionylation conditions (**Supplementary Figure 5**).

To determine the specific residue where O-acetylation occurs, we acquired MS^2^ spectra of the deuteroacetylated mono-acetylated RG-I DP6 (RG-I OAc) at *m/z*=1571 and the non-acetylated RG-I DP6 at *m/z* = 1574 (**Figure 1B**). The target ions (*m/z* = 1571 and 1574) were isolated and subjected to collision-induced dissociation (CID) for fragmentation analysis. The resultant daughter ions and the fragmentation patterns are shown according to the Domon and Costello nomenclature^33^ for RG-I (**Figure 1**). Notably, the *m/z*=773 ion (**Figure 1B**), corresponding to the (B_3_) ion, was present in the MS2 spectra of the deuteroacetylated derivatives of RG-I OAc and RG-I (non-OAc), suggesting that the three residues from the non-reducing end are not O-acetylated. Likewise, the *m/z* = 839 ion (**Figure 1D**) was observed in the MS^2^ spectra of the propionylated derivatives of RG-I OAc and RG-I (non-OAc). The B5 daughter ions at *m/z* 1275 in the deuteroacetylated DP6 RG-I OAc (**Figure 1B**) MS^2^ 1571 and *m/z*=1175 propionylated DP6 RG-I OAc (**Figure 1D**) MS2 1703 suggest that acetylation is present on the reducing end rhamnose. However, *m/z*= 1272 B_5_ ion in the deuteroacetylated DP6 RG-I OAc) (**Figure 1B**) suggest that the acetylation also can be either the second rhamnose or the first galacturonic acid residue from the reducing end.

Further pinpointing the exact position of acetylation was not feasible using MALDI-TOF, due to the instrument’s lack of tandem MS capability and the absence or low abundance of cross-ring fragments in MALDI-TOF/MS/MS spectra of RG-I oligosaccharides. The absence of such fragmentation in MALDI-TOF/MS/MS is primarily attributed to the insufficient sensitivity provided by the instrument, even under CID conditions, to produce these specific fragmentation patterns.

### Finding the O-acetyl position of enzymatically monoacetylated RG-I using direct infusion (DI) ESI-MS^n^

To find the exact position of the acetyl group in RG-I, we used the enzymatically (trichome51 birefringence 3-O acetyl transferase enzyme)^34^ partially (mono) acetylated RG-I DP6 mucilage. The position of the O-acetyl group in the native RG-I (underivatized sample) was also confirmed by nuclear magnetic resonance spectroscopy (NMR) prior to MS analysis. The 2D ^1^H-^1^H correlation spectroscopy (COSY) and heteronuclear single quantum coherence (HSQC) spectra obtained on the native monoacetylated RG-I DP6 (**Supplementary Figure 6 and Supplementary Table 2**) showed the RG-I is acetylated on the 3-O position in the rhamnose, which is also consistent with reported literature chemical shifts.^34^ Then, the partially acetylated RG-I was derivatized (deuteroacetylated or propionylated) and further analyzed using electrospray ionization mass spectroscopy (ESI MS).

ESI-MS allows tandem mass spectroscopy with higher order MS and is equipped with different fragmentation modes, such as collision-induced dissociation (CID), higher-energy collisional dissociation (HCD), ultraviolet photodissociation (UVPD), and electronic excitation dissociation (EED). This makes ESI MS the ideal instrumentation for structural analysis of RG-I oligosaccharides.

ESI MS relies on the efficient desolvation of analytes to generate gas-phase ions. In this process, a high electric field disperses the analyte solution into fine microdroplets. As the solvent evaporates, ions are released and introduced into the mass spectrometer for analysis. Successful ionization depends not only on desolvation but also on the analyte’s solubility in the solvent system. After derivatization with deuteroacetyl and propionyl groups, the sample became more hydrophobic and showed less solubility in methanol (50-90%), which is the most common ESI-MS solvent. Therefore, we used 100% acetonitrile, which is a polar aprotic solvent with low hydrogen-bonding capability as the solvent to solubilize the derivatized RG-I samples for ESI-MS analysis.

For the direct infusion (DI) ESI MS analysis, we employed both collision-induced dissociation (CID) and higher-energy collision dissociation (HCD) MS^n^ at several collision energies to produce higher quality MS^n^ spectra for improved fragment identification. In agreement with prior findings,^35^ we observed that CID resulted mostly in glycosidic cleavages;^35^ fragmentation across the sugar rings (cross-ring fragmentation) was less compared to HCD. This is because cross-ring cleavage requires the simultaneous breakage of two covalent bonds, which is less favorable under CID conditions. In the ion trap, CID typically involves a longer series of low energy collisions with the inert gas, and these often favor low-energy fragmentation pathways (glycosidic cleavages), reducing the detection of labile cross-ring fragments.

In contrast, HCD involves higher energy collisions in a short time frame in ion trap systems. This higher energy imparts more internal energy to the precursor ions, increasing the probability of wider range of sequential fragments and increases the chances of forming more cross-ring fragments. Therefore, we generated MS^1^ and MS^2^ parent ions using CID with a wide isolation window to enrich glycosidic cleavage patterns but acquired MS^3^ spectra using HCD to enhance the detection of cross-ring fragments.

#### Deuteroacetylation

The positive ion mode MS spectrum of deuteroacetylated RG-I DP6 confirmed that the sample consists of derivatized (deuteroacetylated) mixture of mono-acetylated RG-I DP6 (m*/z* 1571), di-acetylated RG-I DP6 (*m/z*=1568), and tri-acetylated RG-I DP6 (*m/z* 1565) in positive sodiated ion forms [M + Na] ^+^ (**Figure 2A**). To further investigate the fragmentation behavior, we performed ESI-MS^2^ on the mono-acetylated RG-I DP6 (*m/z*=1571). CID fragmentation of the *m/z*=1571 parent ion generated similar daughter products in ESI-MS spectra (**Figure 2B)** cleavages labeled in blue) as in MALDI-TOF MS^2^ (**Figure 1B**). These products included ions at *m/z* 1087, 791, 773, 525 and 507, which correspond to Y_4_, C_3_, B_3_, B_3_Y_5_ and B_2_ respectively. The mass of these fragments indicated that the acetyl group was on one of the 3 residues closest to the reducing end of the hexasaccharide. Conversely, the B_5_ and Y_2_ cleavages each yielded a set of two ions that were 3 mass units apart (i.e. the difference between an acetyl and a deuteroacetyl group), suggesting the presence of 2 isomers with an acetyl group in different monosaccharide residues. The B_5_ cleavage gave ions at *m/z* 1272 and 1275, and the Y_2_ cleavage gave ions at *m/z* 585 and 588. The B5 ion with *m/z* 1275 was consistent with a fully deuteroacetylated fragment including the 5 residues from the non-reducing end of Structure E (**Figure 2D**) and thus proving that the acetyl group was located on the reducing end residue; conversely, the B_5_ ion at *m/z* 1272 showed the presence of a different parent ion structure (Structure F) that had the acetylation on a sugar other than the reducing end residue. The Y_2_ ion at *m/z*= 588 was consistent with a fully deuteroacetylated fragment including the two residues closest to the reducing end and thus demonstrated that the acetyl group in Structure F was on the second rhamnose residue from the reducing end. Based on the peak intensity ratios of the *m/z*= 1275 and *m/z*= 1272 B_5_ ion of structure E and F respectively, the sample has 63% of E structure and 37 % of Structure F.

**Figure 2:**
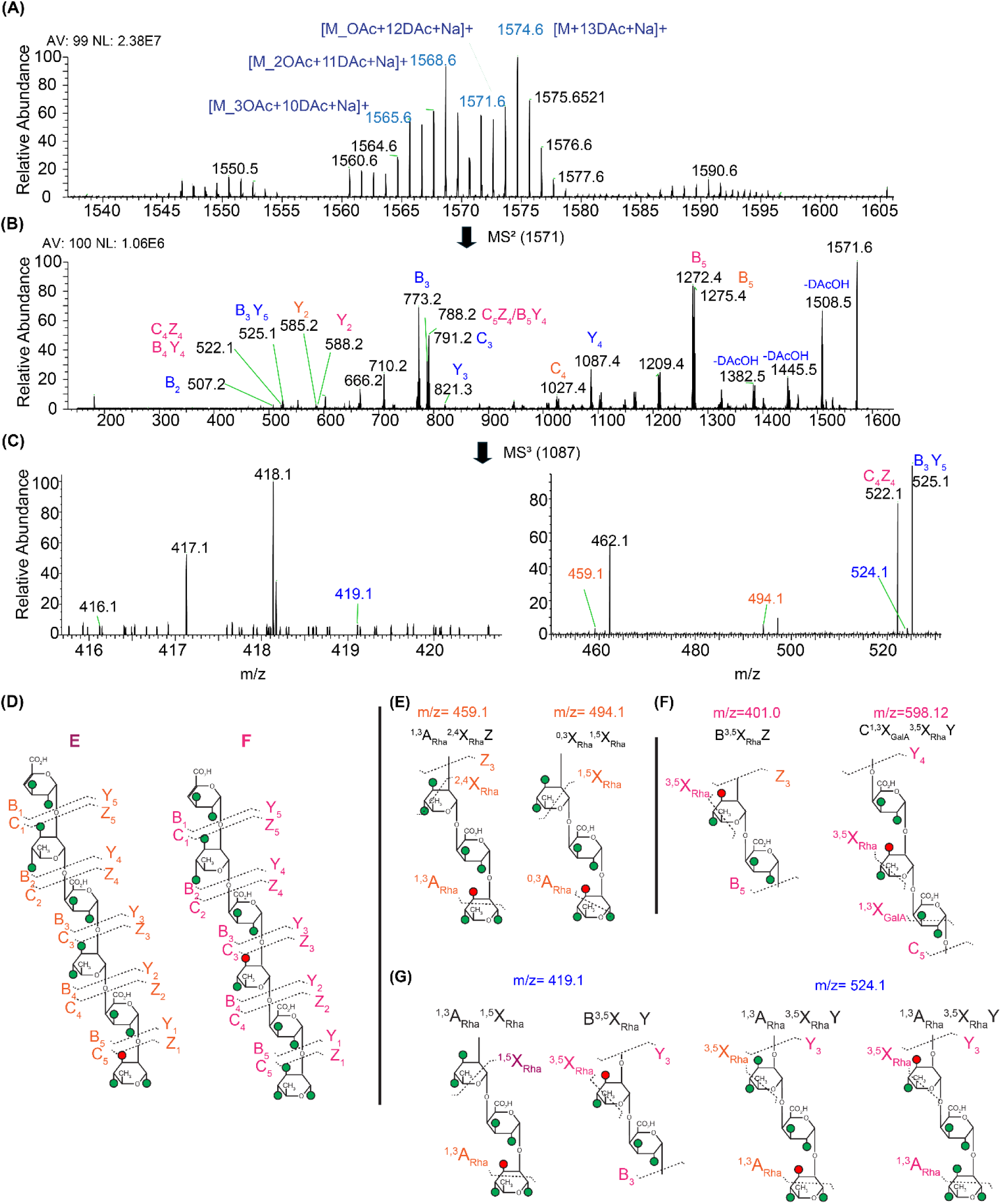
**(A)** Positive ion mode ESI mass spectra of the deuteroacetylated RG-I DP6: **(B)** MS^2^ spectrum of the deuteroacetylated mono acetylated RG-I [M_OAc+12DAc+Na]^+^ ion (*m/z*=1571); DAc denotes deuteroacetyl group. **(C)** Zoomed in MS^3^ spectrum of *m/z*=1087 to show the diagnostic cross ring fragments **(D)** Two possible acetyl position deduced from the MS^n^ spectra structure E and Structure F **(E)** Selected diagnostic cross ring cleavages of structure E *m/z*= 459, and 482 **(F)** Selected cross ring fragments for structure F *m/z* =401 and 598. **(G)** The cross-ring fragments observed for both structure E and F *m/z* 419 and 524. The spectra were obtained in positive mode. The fragments correspond to Structure E labeled in purple, and for Structure F labeled in pink, common fragments for both E and F are labeled in blue. NL: normalized level, AV: averaged number of scans. Non-labeled peaks are not assigned.

We performed further MS^3^ and MS^4^ experiments to provide additional structural information such as linkages and the position of the acetyl group in the rhamnose residue (O-3 or O-4). In the MS³ experiment, we selected the ion at *m/z*=1087 (Y_4_) as the precursor ion and subjected it to HCD MS^3^ fragmentation (**Figure 2C**). This process generated diagnostic cross-ring cleavages that are specifically indicative of the position of the acetyl group, either on O-3 or O-4 of the rhamnose residue. The key fragmentation patterns used for this analysis included ^1,3^A or ^0,3^X, or ^3,5^X cross ring cleavages where cleavage occurs between C_3_ and C_4_ bond of the rhamnose ring (**Supplementary Figure 7**). The *m/z* mass list was pulled from the spectra and each of the theoretical diagnostic cross-ring fragments obtained from Glyco-Workbench was compared with the masses in the spectra to find out the diagnostic cross-ring fragments.

The MS3 process yielded fragment ions at *m/z*=415, 431, 459, 482, 494, and 477, which correspond to ^1,3^A_Rha_^0,2^X_Rha_Z, ^1,3^A ^2,5^X_Rha_Z, ^1,3^A ^2,4^X_Rha_Z_3_, ^3,5^X_Rha_Y_2_, ^0,3^X ^1,5^X_Rha_, and ^1,3^A ^2,4^X Y cleavages respectively (**Table 2**). As shown in **Figure 2E**, fragments at *m/z*= 459 and 482, the ^1,3^A_Rha_ and ^3,5^X_Rha_ cleavages are consistent with 3-O acetylation on the reducing end rhamnose, i.e. Structure E (**Figure 2E**), whereas fragments at *m/z*=401 and m/z 598 show the cleavages of B_5_^3,5^X_Rha_Z_3_, and C_5_ ^1,3^X_Rha_ ^3,5^X_Rha_Y_4_, which are consistent with acetylation is on the 3-O position of the second rhamnose, i.e. Structure F (**Figure 2F**, **Table 2**). Additional supporting evidence came from the molecular ion at *m/z*= 419, and 524 which is giving cleavages for both Structures E and F showing 3-O-acetylation on the rhamnose residue (**Figure 2G**).

**Table 2:** Summarized diagnostic cross ring cleavages for 3-O acetylation observed in the deuteroacetylated monoacetylated RG-I DP6 MS3 spectra *m/z*= 1087 [M+Na]+ for Structure E and Structure F. Shaded are the common cross ring fragments that are observed for both structure E and F.

| Diagnostic cross Ring in MS2 (1571)-> MS3 (1087) |  |  |  |  |  |  |  |
| --- | --- | --- | --- | --- | --- | --- | --- |
| For structure E |  |  |  | For structure F |  |  |  |
| Mass observed | Theoretical | Mass difference | Fragmentation | Mass observed | Theoretical | Mass difference | Fragmentation |
| 415.1271 | 415.1118 | 0.0152 | $^{1,3}A_{Rha}^{0,2}X_{Rha}Z$ | 401.0339 | 401.0961 | 0.0622 | $B^{3,5}X_{Rha}Z$ |
| 431.1141 | 431.1067 | 0.0074 | $^{1,3}A_{Rha}^{2,5}X_{Rha}Z$ | 359.0167 | 359.0856 | 0.0689 | $^{0,2}A_{GalA}^{3,5}X_{Rha}Y$ |
| 459.1397 | 459.138 | 0.0017 | $^{1,3}A_{Rha}^{2,4}X_{Rha}Z$ | 598.1209 | 598.1576 | 0.0367 | $C^{1,3}X_{GalA}^{3,5}X_{Rha}Y$ |
| 482.1717 | 482.1467 | 0.025 | $^{3,5}X_{Rha}Y$ | 627.2243 | 627.1921 | 0.0322 | $^{0,2}A_{GalA}^{0,3}X_{Rha}Y$ |
| 494.1336 | 494.1467 | 0.0131 | $^{0,3}X_{Rha}^{1,5}X_{Rha}$ | 669.2688 | 669.2027 | 0.0661 | $B^{0,3}X_{Rha}Y$ |
| 477.1486 | 477.1757 | 0.0271 | $^{1,3}A_{Rha}^{2,14}X_{Rha}Y$ | 687.260 | 687.2132 | 0.0468 | $C^{0,3}X_{Rha}Y$ |
| 419.1136 | 419.1067 | 0.0069 | $^{1,3}A_{Rha}^{1,5}X_{Rha}$ | 419.1136 | 419.1067 | 0.0069 | $B^{3,5}X_{Rha}Y$ |
| 524.1692 | 524.1572 | 0.012 | $^{1,3}A_{Rha}^{3,5}X_{Rh}Y$ | 524.1692 | 524.1572 | 0.012 | $^{1,3}A_{Rha}^{3,5}X_{Rh}Y$ |

This method also showed the possibility of observing two simultaneous cross ring cleavages from HCD fragmentation. This phenomenon has previously been reported for permethylated oligomers. In these cases, permethylated oligomers yield stable cationic adducts, creating a favorable environment for efficient energy transfer during the collision. This in turn promotes directed fragmentation increasing the likelihood of observing multiple cross-ring cleavages in a single HCD event.^36^

We also evaluated negative ion mode ESI-MS^n^ on the deuteroacetylated RG-I_OAc DP6 sample for its ability to provide structural information on derivatized RG-I (**Supplementary Figure 8**). The negative-ion MS mass spectra showed [M-H]^-^, anionic molecular ion peaks *m/z* 1550, 1547, 1544 for non-acetylated, monoacetylated, and diacetylated RG-I DP6. The MS^2^ fragmentation of the *m/z*=1547 monoacetylated RG-I [M-H]^-^ parent ion produced Y_5_, Z_5_, Y_4_, and Y_3_ fragments at *m/z*=1299, 1281, 1045 and 797 respectively. The negative ion mode analysis also shows the two acetyl positions: at the reducing end and on the second rhamnose. These observations are consistent with the expected structure of the RG-I and align well with the findings from the positive ion mode analysis (**Figure 2**). However, MS^3^ of the *m/z*=1045 precursor ion generated MS^3^ spectra (**Supplementary Figure 8**) with low sensitivity and poor signal-to-noise ratio, with ion signals barely distinguishable from background noise even at reduced HCD collision energies. Therefore, the negative ion mode spectra did not reveal significant information on the acetyl group position. Thus, the negative mode ionization of RG-I ESI-MS^n^ needs further optimization to enhance its effectiveness for structural analysis of RG-I.

#### Propionylation

Having shown that deuteroacetylation is a reliable method to determine the location of native acetyl groups in RG-I, we also wanted to test if propionylation, an alternative derivatization, would provide equal or better results. Propionylation introduces a larger mass shift than deuteroacetylation, which improves spectral separation and may also enhance ionization efficiency in mass spectrometry. Additionally, it avoids complications associated with deuterium exchange, potentially resulting in cleaner and more interpretable spectra. For this purpose, we fully pronionylated the partially acetylated RG-I DP6 and analyzed the product using ESI MS^n^ (**Figure 3**). Similarly, this allowed us to determine the acetyl position in the RG-I oligosaccharides. In the MS^1^ spectrum, ions at *m/z*=1717,1703, and 1689 (**Supplementary Figure 9A**), corresponded to the fully derivatized non-acetylated, mono-, and diacetylated RG-I DP6 respectively. The MS^2^ analysis of fully propionylated monoacetylated RG-I DP6 (*m/z*= 1703) (**Supplementary Figure 9B**), the fragmentation pattern was analogous to that observed in deuteroacetylated spectra (**Figure 2**), yielding key daughter ions, Y_4_, B_3_, and B_5_Y_4_ respectively. Further fragmenting the ion at *m/z*= 1175 (MS^3^, **Supplementary Figure 9C)**. This led to the identification of diagnostic cross-ring fragments for Structure E at *m/z* = 499 and 515 (**Supplementary Figure 9D**), where the acetyl group is on the 3-OH position on the reducing end rhamnose. The MS^3^ spectra (**Supplementary Figure 9C**) also showed evidence for Structure F by the *m/z*=153 ion, which is characteristic of a B ^3,5^X Y cross-ring cleavage on the third rhamnose residue (**Supplementary Figure 9E)**.

**Figure 3:**
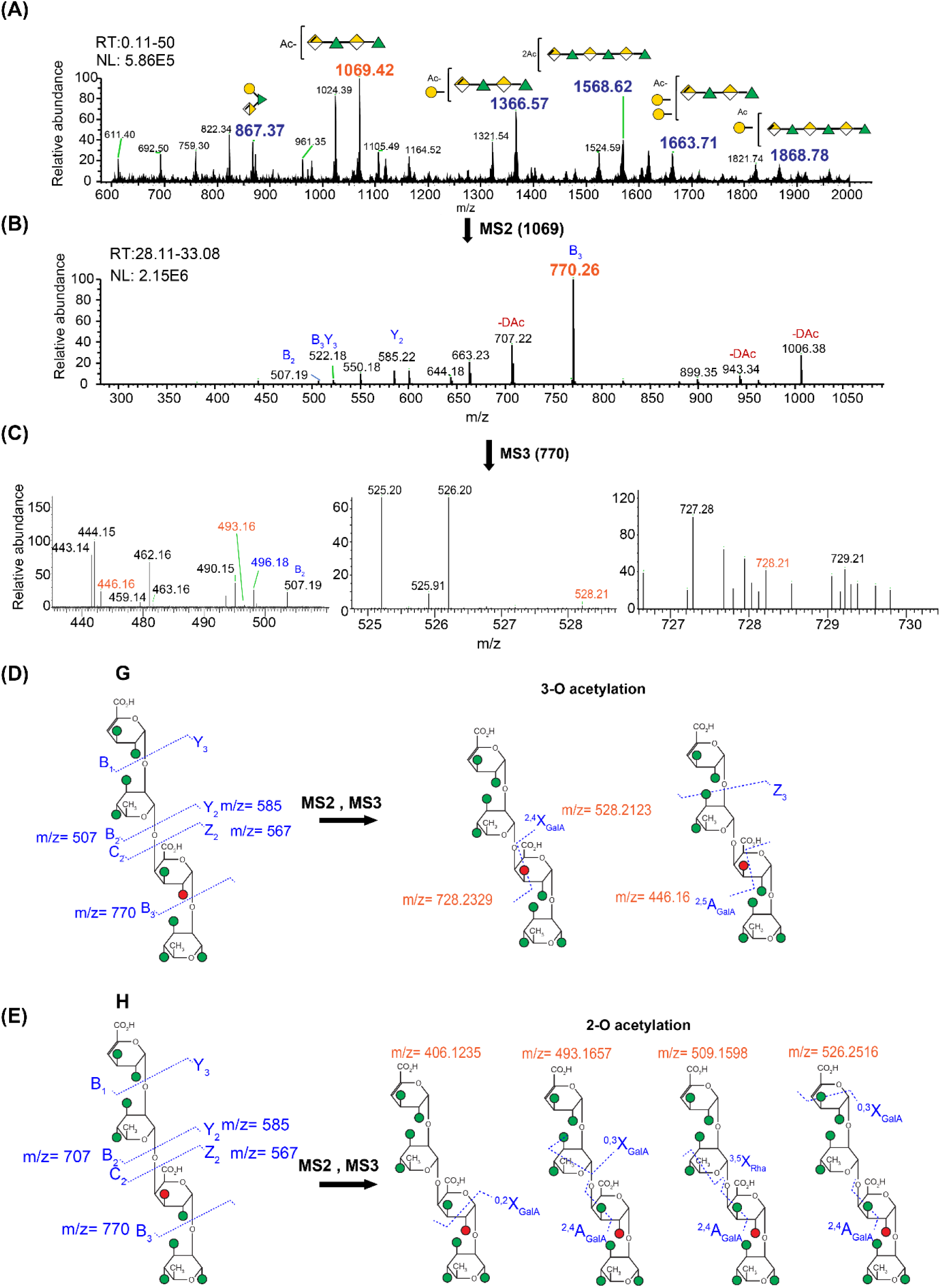
**(A)** Deuteroacetylated RG-I celery LC-ESI MS average spectrum in positive ion mode. The spectrum is the average of the retention time (RT) 0.11 to 50 min. **(B)** Average MS^2^ spectrum (RT 28.11-33.08 min) of the [M–Na] ^+^ ion *m/z*=1069. Symbol nomenclature for glycan (SNFG) shows the identified structures for *m/z* in the MS spectra. The ambiguous linkage positions of acetyl and galactose residues are indicated in brackets. (**C)** The MS^3^ spectrum was obtained from the *m/z*=770 fragment ion. The spectrum was average from 0.3-50 min elution averaged 1590 number of scans. The two possible structures identified from the fragments observed in the MS^2^ spectrum of the ion at *m/z*=1069 are (**D)** Structure G: RG-I 4DP4 with 2-O acetylation in the second GalA residue and (**E**) Structure H: RG-I 4DP4 with 3-O acetylation in the second GalA residue. Also, the diagnostic cross-ring fragmentation patterns and fragments observed for Strucure G and H. LC elution profiles are shown in the Supplementary Figure 11

Both deuteroacetylation and propionylation proved to be effective derivatization methods for determining the acetyl position in the RG-I oligomers, providing complementary fragmentation patterns that reinforced the structural assignment of the acetyl position. Though it was successful for RG-I DP6, the use of a bulkier acyl group for DP oligomers (not tested) could potentially reduce solubility due to increased hydrophobicity. Overall, deuteroacetylation is the preferred method, as it offers better resolution for tracking native O-acetyl modifications, but both approaches can be used together to enhance confidence in the structural assignment.

The results from ESI-MS analysis were consistent with NMR data (**Supplementary Figure 6 and Supplementary Table 2**), both indicating the presence of mono-, di-, and triacetylated species with acetylation on the 3-O position on the rhamnose. While ESI-MS suggested that the monoacetylation could occur either on the reducing-end rhamnose or on the second rhamnose residue, the NMR data - along with previously reported acetylation patterns of TBR enzyme products supported monoacetylation only at the reducing-end rhamnose.^34^ Due to the similarity in chemical shifts and potential peak overlap, the two mono-acetylated isomers could not be clearly distinguished by NMR alone. In contrast, ESI-MS was able to resolve these isomers, highlighting its utility in detecting positional isomers that are difficult to differentiate by NMR.

### Finding the O-acetyl position of celery RG-I using LC-ESI-MS^n^

To apply our method to an isolated sample with native acetylation, we purified RG-I from celery and digested it with RG-I lyase enzyme to generate oligosaccharides. We screened the native digested sample using LC-MS, which showed DP4 and DP6 RG-I oligosaccharides with one or two acetyl groups or single galactose branches as components of the sample (**Supplementary Figure 10**). Although the MS² spectra of the underivatized sample confirmed the presence of RG-I oligomers, they did not permit unambiguous assignment of the acetylated residues. Also, due to the complexity of the mixture and low sample purity, the signals of the target oligomers were difficult to detect, as they were obscured by interfering ions from solvents, buffers, and other co-eluting components.

However, these oligosaccharides represent an excellent test case to demonstrate the power of LC-MS^n^ of deuteroacetylated derivatives. Due to the complexity of the background and the diversity in the RG-I oligomers present in the digested celery RG-I sample, the fully deuteroacetylated RG-I oligomers were not detectable by the direct infusion ESI method. Therefore, we analyzed the deuteroacetylated RG-I oligosaccharide sample using nanoflow liquid chromatography (nanoLC), combined with MS^n^ detection (LC-MS^n^) in positive mode and observed a multitude of sodiated oligomer ions.

The MS profiling of the deuteroacetylated celery RG-I oligomers in LC-MS and MALDI-TOF **(Figure 3A and Supplementary Figure 11**) showed peaks belonging to mono- or diacetylated oligosaccharides derived from four to seven RG-I backbone residues and also shows RG-I oligomers containing galactose side chain residues. For each *m/z* peak, multiple isomers could be constructed that differed in the placement of acetyl group on individual residues and also in the location of the branching in the sequence. This complex mixture of RG-I oligomers showed an LC-MS elution order that correlated with the degree of polymerization, degree of acetylation, the branching pattern and the length of the branching. The use of a C18 reversed-phase column enabled separation based on hydrophobicity, where oligomers with higher acetylation and longer or more branched side chains exhibited longer retention times due to increased interactions with the column matrix (**Supplementary Figure 12**). The molecular ion of *m/z*= 1069 (DP4 RG-I_OAc) eluted first, followed by, *m/z*= 1366 (DP4 RG-I_OAc with one galactose side chain) and *m/z*= 1568 (DP6 RG-I_2OAc). These ions were chosen for further analysis of acetylation position by tandem MS (**Figure 3A**).

#### m/z= 1069 – deuteroacetylated tetrasaccharide RG-I_OAc tandem MS analysis

We selected the perdeuteroacetylated RG-I DP4 backbone with a single acetyl group (*m/z* = 1069) as the precursor ion and analyzed it by MS² using CID. The fragment ions obtained [*m/z*=770 (B3), 585 (Y2), 507 (B2)] indicated that the acetyl group was primarily located on the GalA residue (**Figure 3B**) rather than on the reducing-end rhamnose [*m/z*=773 (B3)], which accounted for less than 2% of the total population (**Supplementary Figure 13**).

Further MS³ analysis of the *m/z* = 770 (B3) fragment ion using HCD generated both glycosidic and cross-ring fragments (**Figure 3C**). Comparison of the observed fragment ions with theoretical fragments generated using GlycoWorkbench identified diagnostic cross-ring ions at m/z = 446, 528, 728, and 409, corresponding to ^2,5^A_GalA_Z, ^2,4^X_GalA_, ^0,2^A_GalA_, ^0,2^ X_GalA_ fragments (**Supplementary Table 3**). These fragments confirmed that the major site of acetylation was at the 3-O position of the GalA residue (**Figure 3D**). In addition, minor diagnostic fragments consistent with 2-O-acetylation were also detected, indicating a small proportion (∼14%) of 2-O-acetylated GalA (**Figure 3E**). Overall, the RG-I DP4 backbone was predominantly 3-O-acetylated on the GalA residue (approximately 86%) (**Supplementary Figure 14**).

#### m/z= 1366 deuteroacetylated branched pentasaccharide RG-I_OAc tandem MS analysis

The addition of branching significantly increases the complexity of the acetylated RG-I. The molecular ion *m/z*=1366; which comes from an RG-I pentasaccharide comprising a DP4 RG-I backbone with a single galactose branch and one acetyl group. The Gal residue can be either on the reducing end Rha (Structures J, K, and L in **Figures 4C and Supplementary Figure 15**) or the Rha next to the non-reducing end (Structures M, N, and O in **Supplementary Figure 15**). Together with the various possible positions of the acetyl group, this gives rise to multiple isomers.

**Figure 4:**
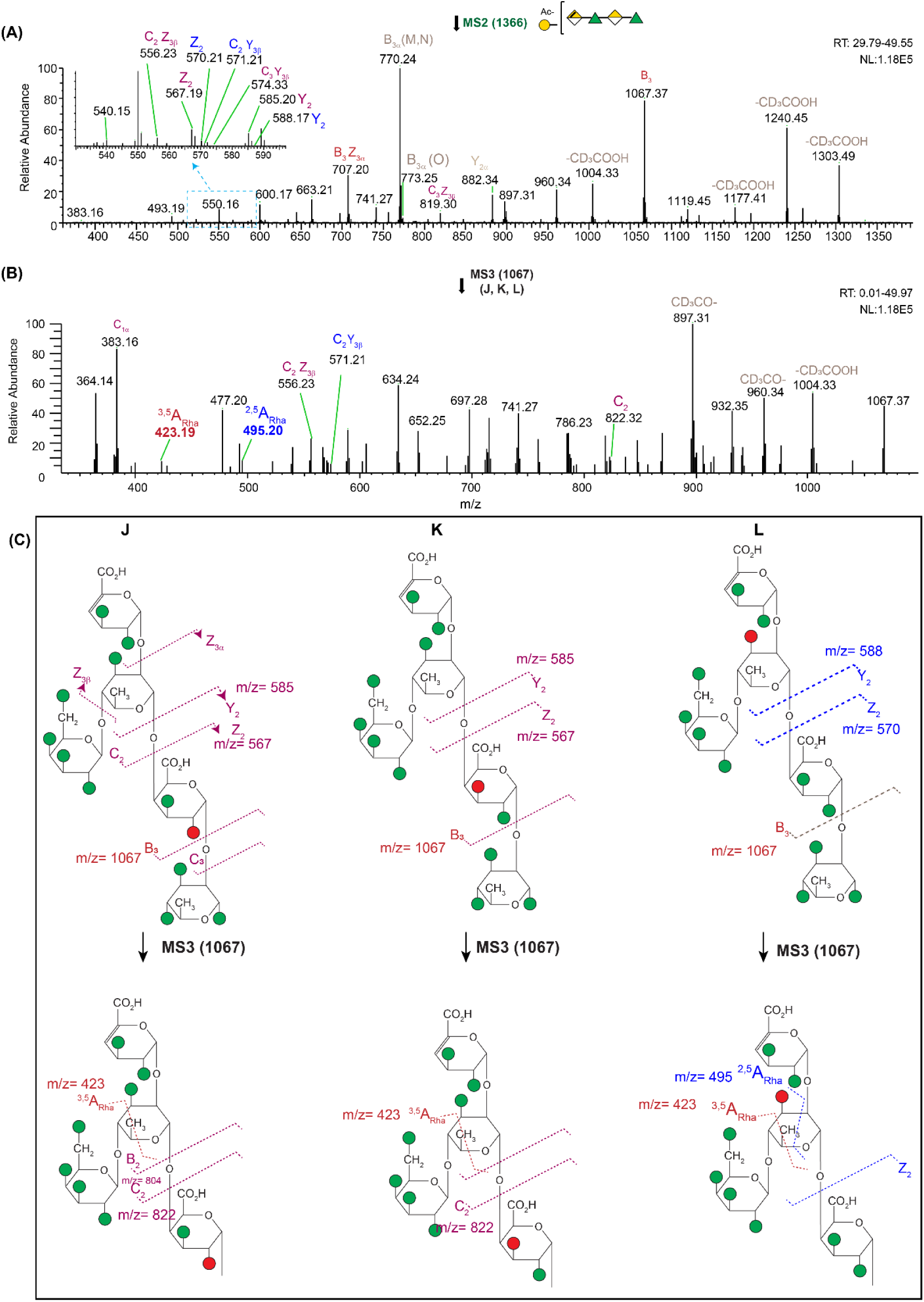
**(A)** MS^2^ spectrum of *m/z*=1366 [M+Na]+ inset shows the zoom in region marked n blue box. The spectra is average from RT 29.79-49.55 min **(B)** MS^3^ spectrum of *m/z*= 1067 for J, K, L isomers average from RT 0.01 49.97 min **(C**) The J, K L structures and cleavages observed in the MS spectra are labeled. The red circle denotes the acetyl groups, and the green circles are deuteroacetylated groups. The *m/z* 423 diagnostic cross ring cleavage present in J, K and L confirms that the rhamnose is branched at the 4-O position. NL: normalized intensity level. The structures of the non-labeled peaks are not determined.

To narrow down the possible isomers and to get more structural information, we isolated the molecular ion at *m/z*=1366 and subjected it to MS^2^ analysis (**Figure 4A**). The resulting spectrum revealed fragment ions that provided critical insight. The fragment at *m/z*=1067 (B3 cleavage) was consistent with the structure of J, K, and L. Conversely, the fragment at *m/z*= 770 (B_3α_) was only consistent with the M and N isomers, indicating 2-O or 3-O acetylation on the GalA residue (**Supplementary Table 3 and Supplementary Figure 16C)**. Further diagnostic ions at *m/z* = 585, 556, 567, and 574 (Y_2_, C_2_Z_3β_, Z_2_, and C_2_Y_3β_) were observed exclusively in the J and K isomers (**Figure 4C and Supplementary Table 4**). Ions at *m/z* = 588, 571, and 570 (Y_2_, C_3_Y_3β,_ and Z2) were specific only to the L isomer (**Figure 4C**: Structure L), and *m/z* 773 and 616 were for the O isomer, suggesting the presence of the 3-O-acetylation on the branching rhamnose (**Supplementary Table 3**). The MS^2^ spectrum showed fragments indicative of acetylation on both GalA and Rha residues. These isomers are co-eluting together and cannot be separated chromatographically (**Supplementary Figure 17**).

To gain a deeper understanding of the specific isomers present (J, K, and L isomers), we performed MS^3^ analysis on the *m/z* =1067 B3 fragment (J, K, and L **Figure 4B** and **Supplementary Table 5**). The MS^3^ spectrum confirmed the presence of acetyl groups on both GalA and Rha, further supporting the identification of the J, K and L isomers. A fragment at *m/z*= 423.19 confirmed the galactose branching on the 4-O position of rhamnose, corresponding to a ^3,5^A_Rha_ cleavage for J, K and L isomers (**Figure 4C** and **Supplementary Table 5)**. These results support the conclusion that the second rhamnose residue is linked as → 2,4)-α-L-Rha-(1→, leaving only the 3-OH position free.

The branching galactose at the 4-O position having been established, the fragments at *m/z* = 495, 507, 668, 717, and 745 (corresponding to ^2,5^A_Rha_, ^1,5^A_Rha_Z, ^3,5^A_GalA_Z, B^1,3^X_Rha_ and^1,4^A_Rha_ respectively, see **Figure 4C**) prove that the rhamnose acetylation is present on the 3-O position, confirming Structure L. On other hand, it is challenging to definitively assign 2-O/3-O acetylation in the GalA residue (Structure J and K) with the current MS data due to many possibilities of cleavage pathways and multiple coexisting isomers and fragments.

To determine the location of the O-acetyl group on the GalA residue, we performed MS^3^ of m/z 770 (obtained from 1366 MS^2^) (**Supplementary Figure 16**). The spectra revealed diagnostic peaks for both 3-O-acetylation (M isomer) and 2-O acetylation (N isomer) **(Supplementary Figure 16 B-C and Supplementary Table 5)**.

#### m/z= 1568 – perdeuteroacetylated linear hexasaccharide RG-I_2OAc tandem MS analysis

We also studied perdeuteroacetylated DP6 RG-I with two acetyl groups (*m/z*= 1568) with tandem MS (**Supplementary Figure 18)**. The MS^2^ CID spectrum of the *m/z*=1568 gave daughter ions characteristic of P, Q, R, S and T (**Supplementary Figure 18a, b**). The product ion *m/z*=1084 (Y_4_), ion confirms the P, Q, R, S, and T and *m/z*= 785, 1269 confirms the P and Q structures (**Supplementary Figure 18**). B_3_ (*m/z*= 770) showed that two acetyl groups are like in the structure Q, R, S, and T. Further MS^3^ analysis of *m/z* = 1084 (**Supplementary Figure 18c**) gave diagnostic cross-ring peak *m/z*=461.15 cleavage ^2,5^A_GalA_Y_3_, which shows the presence of 3-O acetylation in GalA residue adjacent to an acetylated rhamnose corresponding to Structure P (**Supplementary Figure 18d**; Structure P). However, due to the complexity of the structure (many possible acetylation positions) and due to limitation in the diagnostic cross-ring cleavages the acetylation position of the rhamnose (P isomer) or other structures cannot be confirmed.

Overall, the structural analysis of perdeuteroacetylated celery RG-I oligomers using the LC-ESI-MS^n^ revealed a highly heterogeneous composition with multiple isomers arising from variations in acetylation and branching. The DP4 RG-I backbone (*m/z* = 1069) was found to carry a single acetyl group, predominantly at the 3-O position of the GalA residue, with minor 2-O acetylation also detected. The acetylated RG-I DP4 with a galactose branch (*m/z* =1366) exhibited acetylation on both GalA and Rha residues, with the galactose branch linked to the 4-O position of Rha. /MS^2^ and MS³ analysis of DP6 RG-I with two acetyl groups (*m/z* = 1568) indicated that acetylation occurred on both GalA and Rha, with some isomers showing adjacent acetylation on these residues. These findings highlight the structural complexity of RG-I, with acetylation and branching patterns contributing to its diverse isomeric forms.

To complement the MS data, we performed NMR analysis on the native RG-I from celery (undigested). This provided an evidence for rhamnose 3-O-acetylation and the presence of branching galactose on the acetylated rhamnose residues (**Supplementary Figure 19**).^37^ However, due to peak overlap and spectral complexity, determining the acetylation of GalA was not possible by NMR. In contrast, the combination of depolymerization, derivatization and ESI-MS^n^ of the oligomers offer a faster and more reliable approach for identifying acetylation sites, it also provides the capability of seeing minor modification (such as 2-O acetylation of GalA) which cannot be observed by NMR. While further purification of digested RG-I oligomers could facilitate acetyl position determination using NMR, its applicability is constrained by the relatively high sample quantity that is required. Thus, MS proves to be a more effective method for pinpointing acetylation sites compared to NMR in the case of RG-I.

## CONCLUSION

In conclusion, this study demonstrates the effectiveness of advanced tandem mass spectrometry approaches, including CID, and HCD for the structural elucidation of RG-I oligomers with distinct O-acetylation patterns. By employing combinatorial fragmentation strategies and leveraging LC-MS separation technique, we enhanced the resolution of complex mixtures, facilitating precise acetyl group localization. The introduction of trideuteroacetyl and propionyl modifications successfully stabilized O-acetyl groups, mitigating challenges associated with acetyl migration and loss during MS^n^ analysis and increase the sensitivity to pinpoint the acetyl position. However, conventional CID and HCD methods primarily generate glycosidic and cross-ring cleavages, sometimes limiting the ability to differentiate between closely related isomers when it comes to complex mixtures. Incorporating ultraviolet photodissociation (UVPD), electronic excitation dissociation (EED) and electron transfer dissociation (ETD) could overcome these limitations by providing enhanced diagnostic cross-ring fragmentation pathways, preserving labile modifications, and offering more comprehensive structural characterization. This approach provides for the first time valuable insights into pectin O-acetylation, with implications for understanding the functional roles of RG-I modifications in plant biology. Furthermore, this strategy can be extended to other acylated oligosaccharides, offering a powerful tool for studying enzymatic regiospecificity and structural diversity in complex polysaccharides.

## DATA AVAILABILITY

All relevant data that support the findings of this study are available within the article and Supplementary Information.

## SUPPLEMENTARY INFORMATION

Additional method details, Supporting Figures for the results and Supporting Table for experimental results are summarized.

## ACKNOWLEGMENTS

This work was supported by the U.S. Department of Energy, Office of Science, Basic Energy Sciences, Chemical Sciences, Geosciences and Biosciences Division, under award DE-SC0015662 and the National Institutes of Health (NIH)-funded R24 grant (R24GM137782) to Parastoo Azadi. This work was supported by GlycoMIP, a National Science Foundation Materials Innovation Platform funded through Cooperative Agreement DMR-1933525. We wish to acknowledge Lubana Shahin, Dr. Liang Zhang, and Dr. Breeanna Urbanowicz for providing the samples used in this study. Thank you to Jiri Vlach and Ian Black for the in valuable discussions and insight to this study.

## AUTHOR CONTRIBUTIONS

L.D.F. prepared the derivatized samples and L.D. F, X.Y, and S. A. conducted MS experiments. L.D.F. X.Y, and S. A analyzed the MS data and interpreted the structural data. P.A and C.H designed and supervised the project. All authors contributed to the manuscript writing and editing.

