## Supplementary Information for "Mass Spectrometric Determination of Site-Specific O-Acetylation in Rhamnogalacturonan-I Oligomers"

**Supplementary Methods 1:**

Summarizing the General method for Determining RG-I Acetylation Sites by Tandem Mass Spectrometry

1. Sample Preparation:
   The partially acetylated oligosaccharides or polysaccharide fragments are isolated and purified. (partial hydrolysis or enzymatic digestion to produce oligomers). Derivatize the oligomers (deuteroacetylation or propionylation).

2. Mass spectroscopy Ionization:
   The sample is introduced into the mass spectrometer using ESI (electrospray ionization) or LC-MS generating molecular ions (either positive or negative ion mode).

3. Selection of Precursor Ion: A specific acetylated molecular ion (e.g., $[M–H]^-$/[M-Na]+) corresponding to the desired degree of polymerization (DP) and acetylation is selected for fragmentation.

4. Fragmentation ($MS^2$ / $MS^n$ Analysis): The precursor ion is subjected to collision-induced dissociation (CID) or higher-energy collisional dissociation (HCD).

5. Interpretation of Fragment Ions:

Glycosidic cleavages (B/Y or C/Z ions) indicate the sequence of monosaccharide residues.

Cross-ring cleavages (A- and X-type ions) provide structural information about substitutions (e.g., acetylation sites) on specific sugar residues.

- Predict the possible structures in Glyco Work bench
- Narrow down the predicted structures by comparing the MS2 daughter ion obtained.
- Predict the possible diagnostic cross-rings based on the acetyl position on the residues using Glyco Work bench
- The *m/z* mass list is pulled from the spectra and each of the theoretical diagnostic cross-ring fragments obtained from Glyco-Workbench was compared with the masses in the spectra to find out the diagnostic cross-ring fragments.

6. Validation and Cross-Verification:
   The proposed acetylation sites are confirmed by comparing experimental spectra with theoretical fragmentation patterns, or complementary techniques such as NMR spectroscopy.

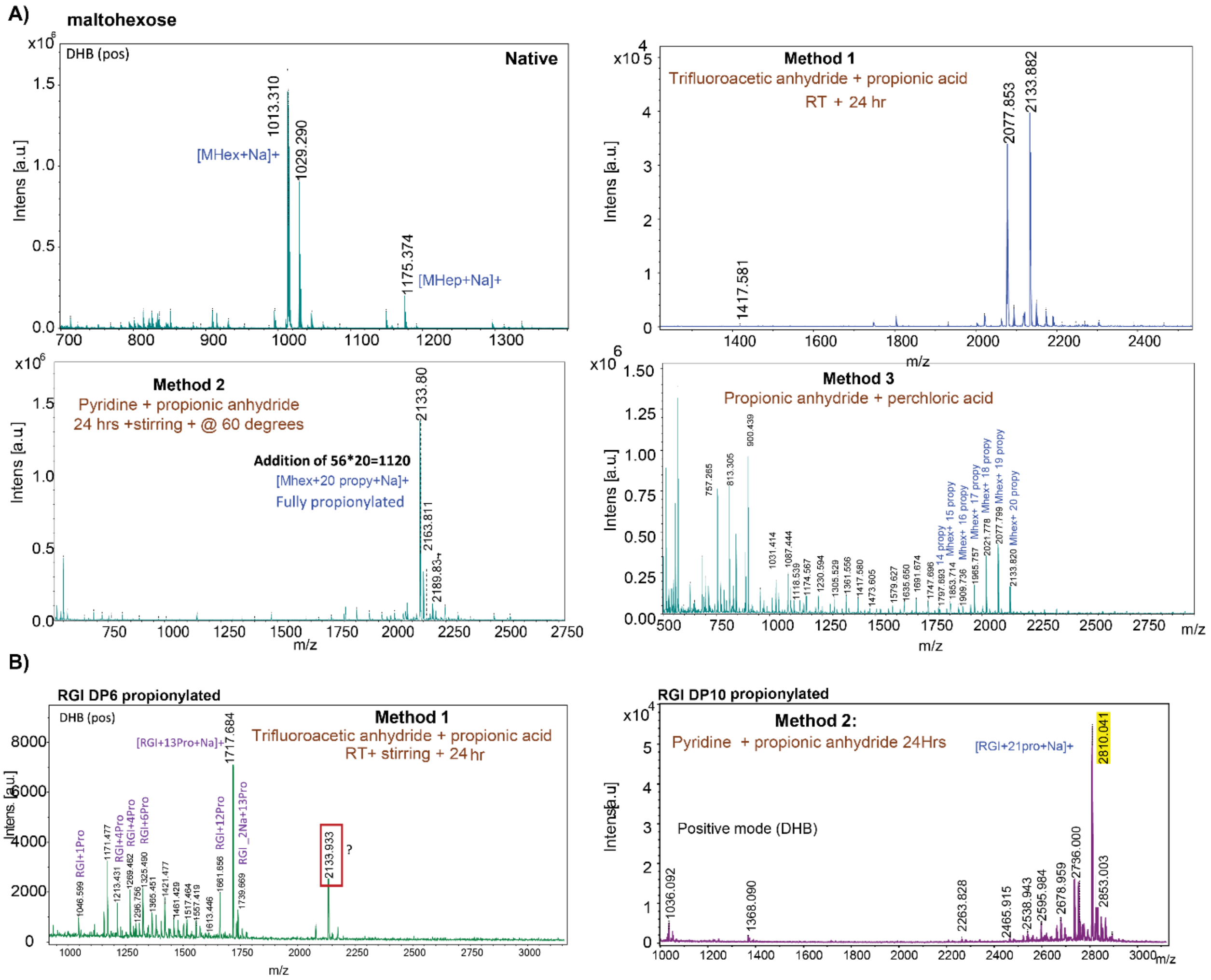


**Supplementary Figure 1:** MALDI-TOF spectra of **A**) Maltohexoes (underivatized), and acylated maltohexose with three different methods; Method **1**- trifluoracetic anhydride + propionic acid, Method **2**- pyridine + propionic anhydride, Method **3**- Propionic anhydride +perchloric acid. **B)** Derivatized RG-I DP6 with Method **1** (left) and Method **2** (right).

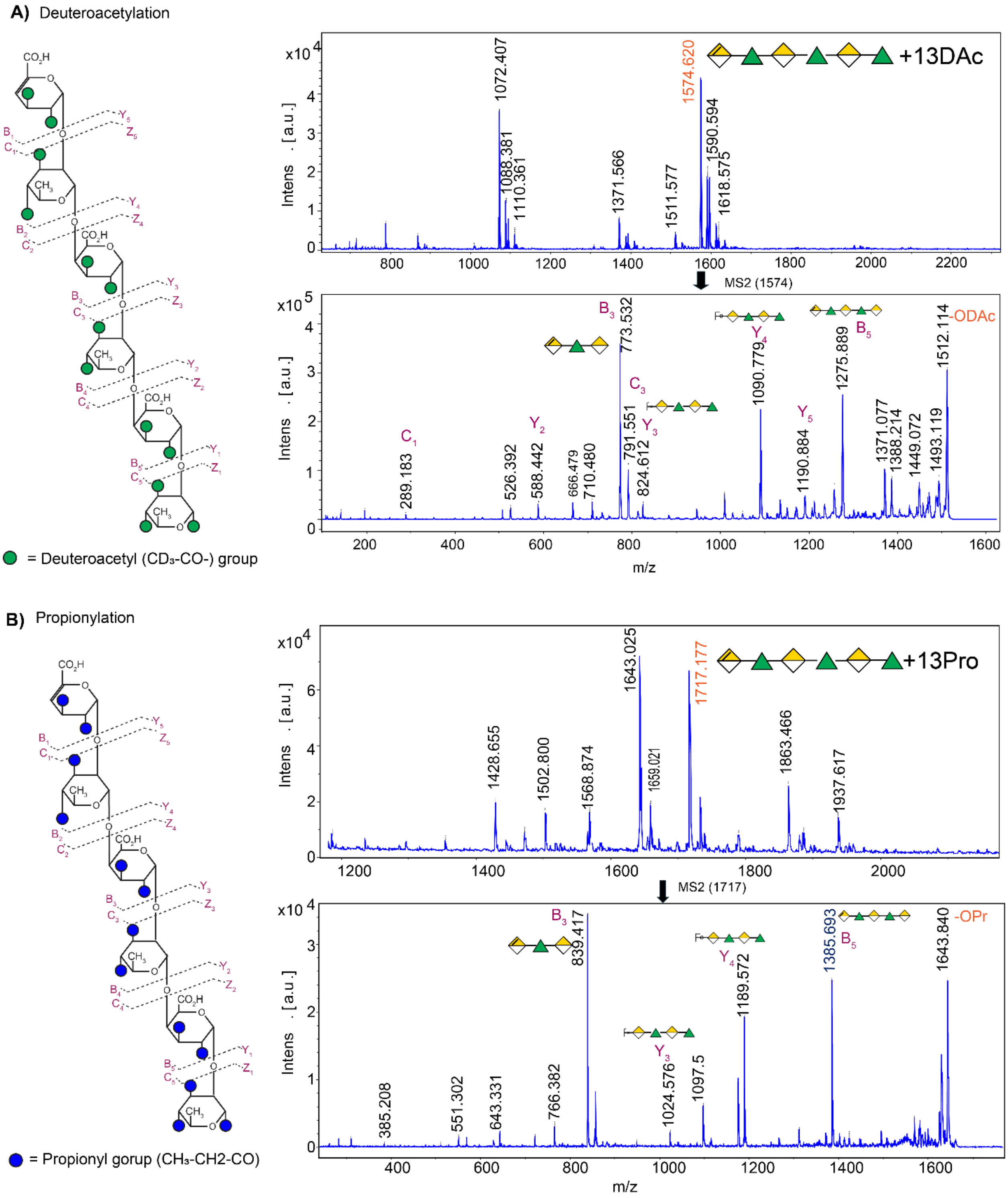


**Supplementary Figure 2:** MALDI-TOF spectra of **A)** Deuteroacetylated RG-I DP6 (m/z 1574 represent fully deuteroacetylated RG-I and the MS$^2$ spectra of the $m/z$ = 1574 parent ion. **B)** Propionylated RG-I DP6 and the MS2 spectra of the $m/z$ = 1717 parent ion. -DAc denotes deter acetyl groups and “-Pro” denotes propionyl groups.

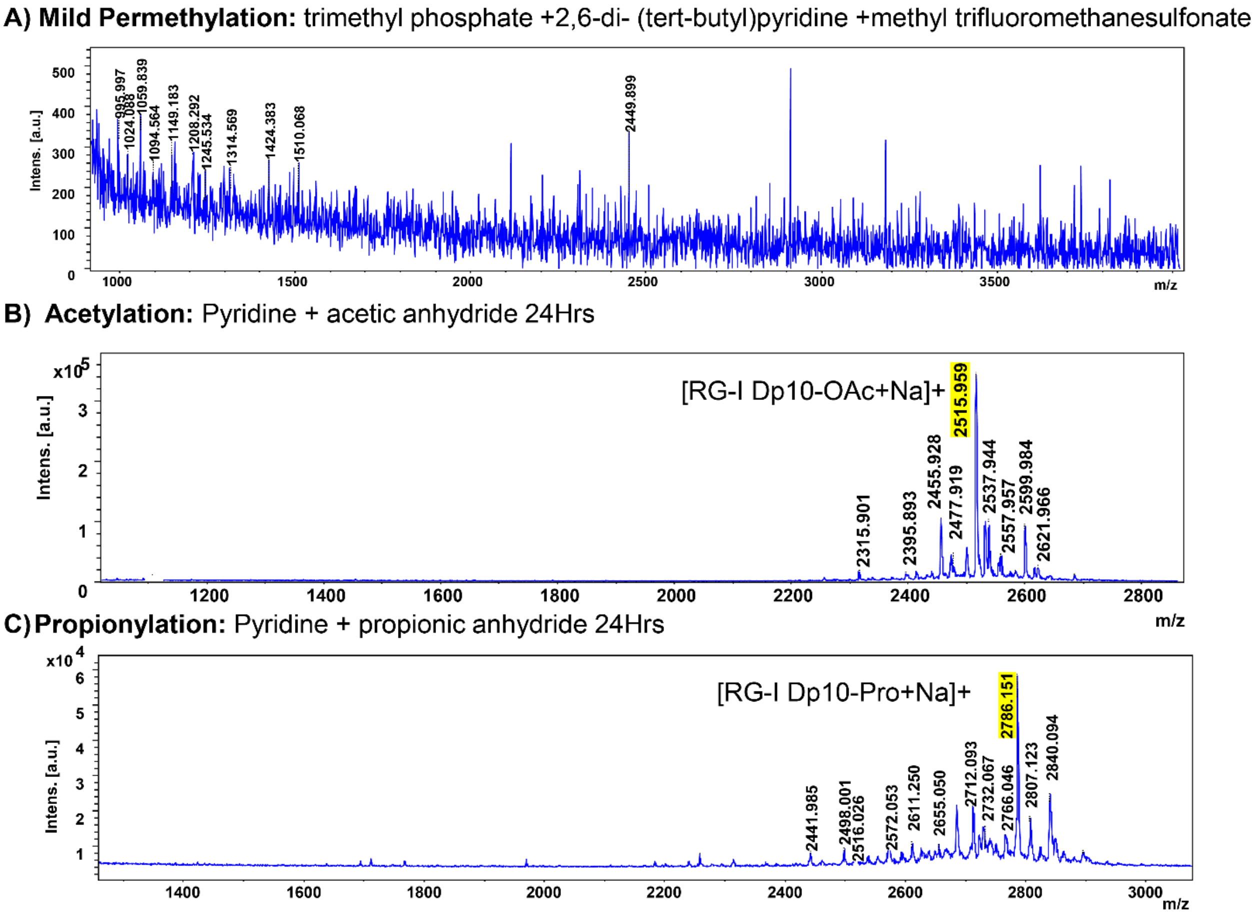


**Supplementary Figure 3:** MALDI-TOF spectra of a) mild permethylated RG-I DP6 b) acetylated RG-I DP 10 and c) Propionylated RG-I DP10.

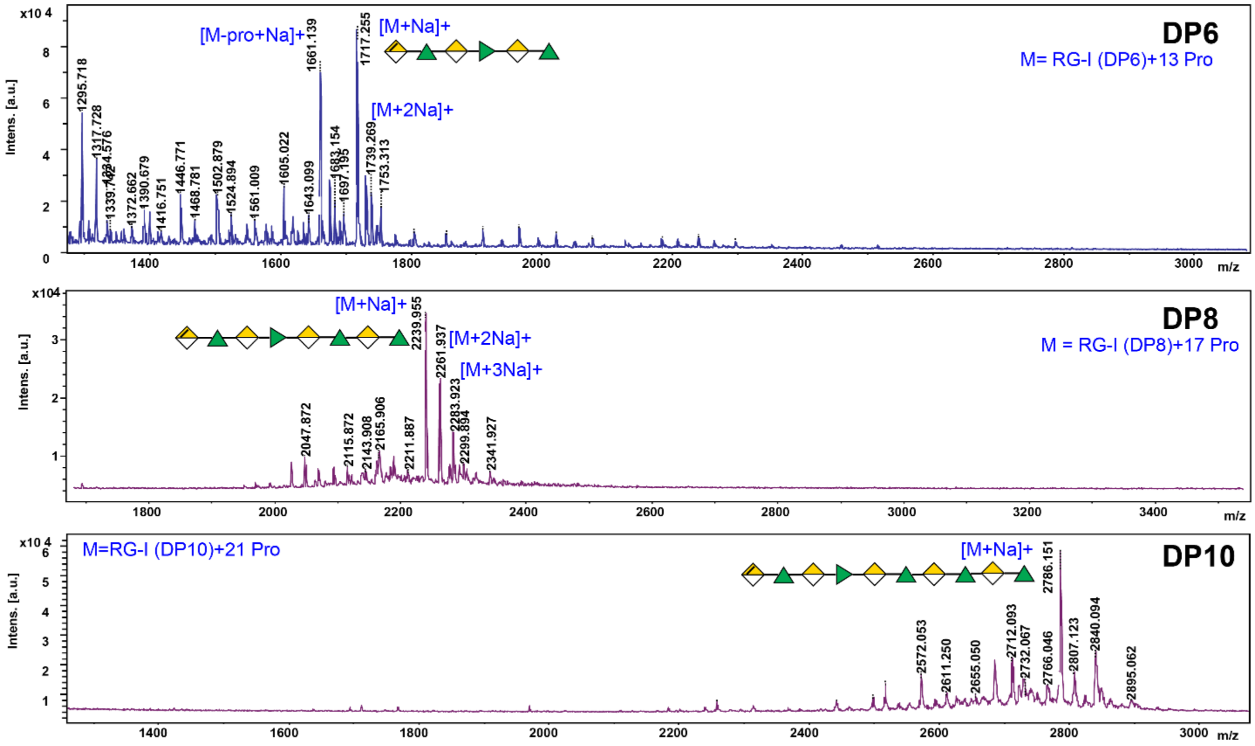


**Supplementary Figure 4:** MALDI-TOF spectra of propionylated DP6, DP8 and DP10 RG-I. Pro denotes propionyl groups.

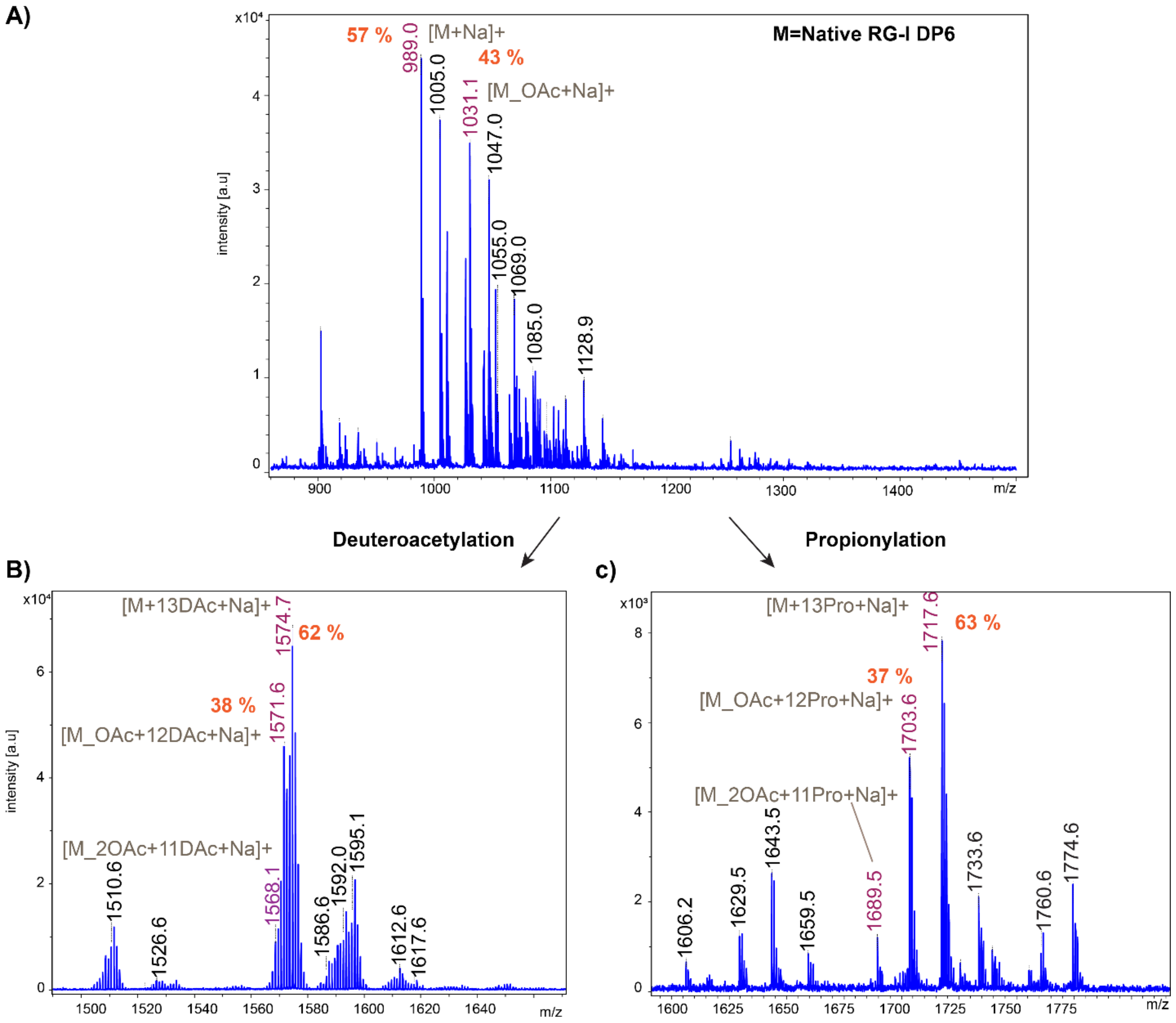


**Supplementary Figure 5:** Enzymatically mono acetylated RG-I DP 6 oligomer MALDI-TOF MS spectra in positive ion mode $[M+Na]^+$. The ratio between non-acetylated to mono acetylated RG-I DP6 remain same after derivatization a) MALDI-TOF MS spectra of the native underivatized oligomer shows a mixture of non-acetylated RG-I (m/z =989; 57%) and mono acetylated RG-I DP6 (m/z=1031;43%), minor amount of di acetylated RG-I DP (m/z=1073) b) Fully deuteroacetylated RG-I DP 6 MALDI-TOF MS spectra shows ratios between non-acetylated RG-I m/z=1574 (62%) and mono acetylated RG-I m/z=1571 (38%). c) MADLI-TOF MS spectra of RG-I DP6 after propionylation. Similarly, the ratio between non-acetylated to acetylated RG-I DP6 remains same (63% : 37%). Pro denotes propionyl groups and Dac denotes deuteroacetyl groups.

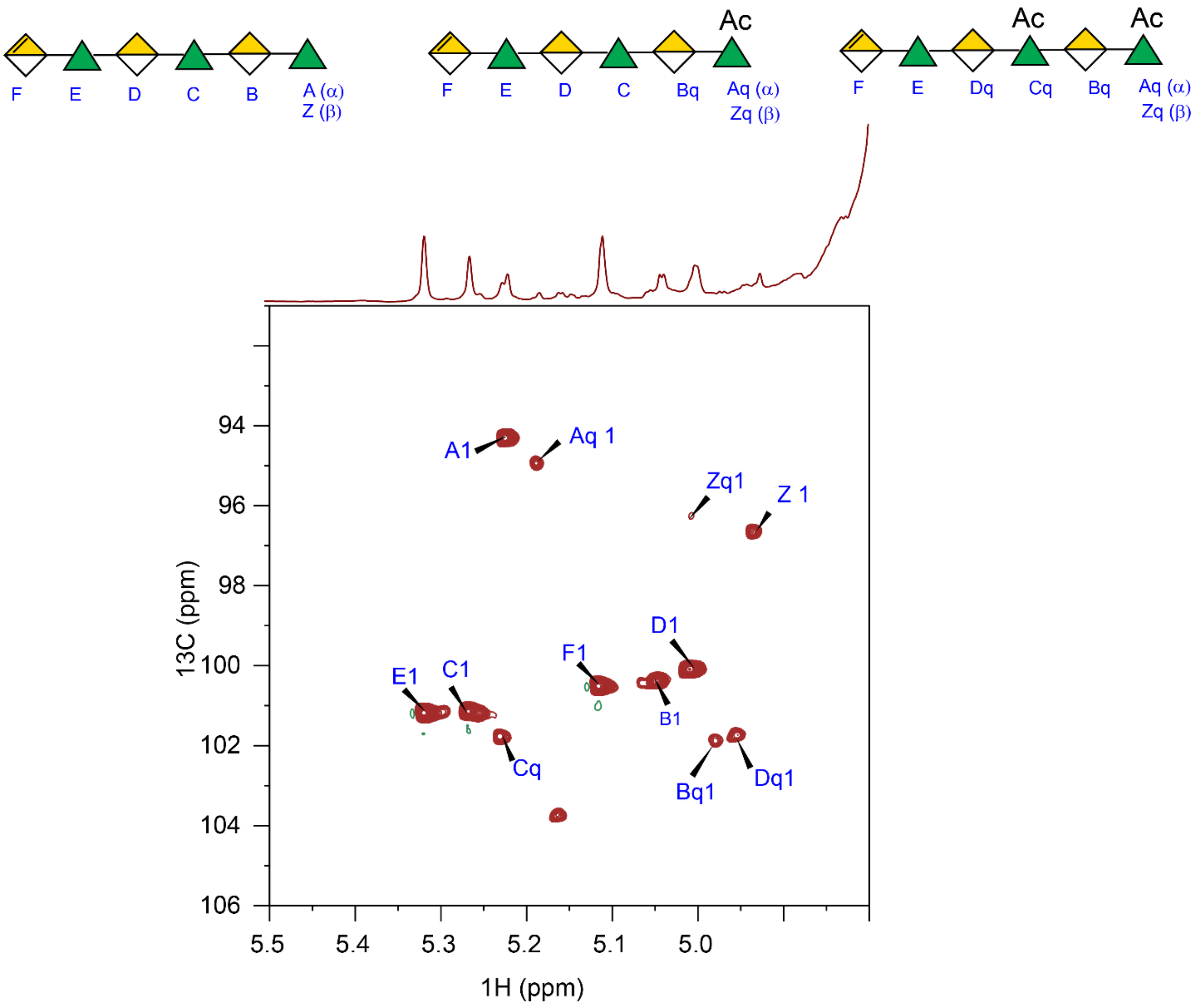


**Supplementary Figure 6:** 2D $^{1}H$-$^{13}C$ HSQC spectra of the partially acetylated RG-I DP6. Mixture of partially acetylated RG-I DP6 present in the sample (top). The 2D HSQC spectra shows the diagnostic anomeric peaks for the non-acetylated RG-I DP6, mono acetylated and di acetylated RG-I DP6. The HSQC also confirms the 3-O acetylation in the rhamnose residue in the RG-I.

.

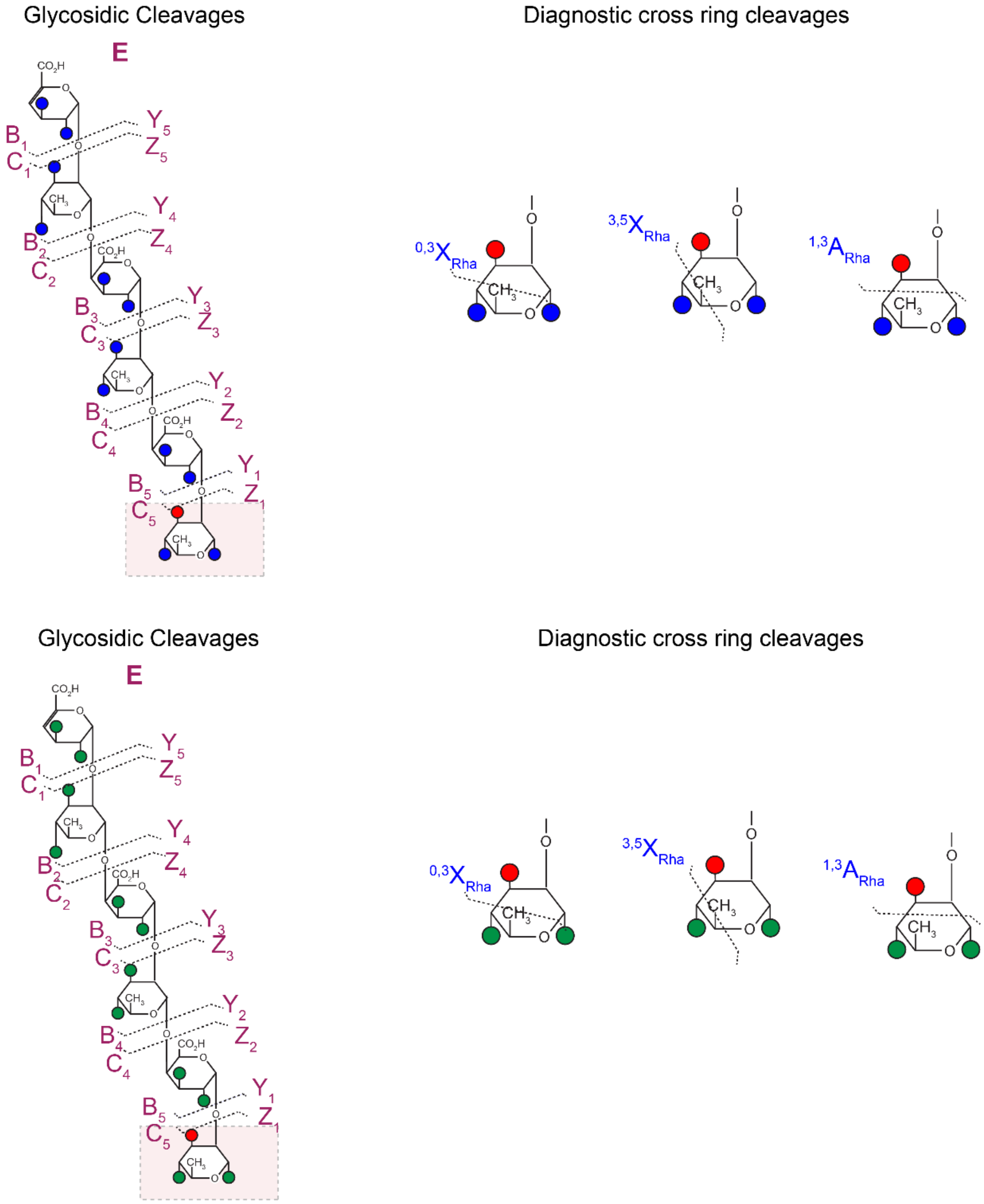


**Supplementary Figure 7:** The diagnostic cross-ring cleavages (A and X cleavages) for DP6 RG-I_OAc. The Fragmentation nomenclature according to Domon and Costello. In order find the position of the acetyl group the cross-ring cleavage needs to happen in the covalent bond between two carbons connected to the acetyl group and the free hydroxyl group. Blue circle denotes the free hydroxyl with derivatization with propionyl groups and green circle denotes the deuteroacetylation, red circle: acetyl group.

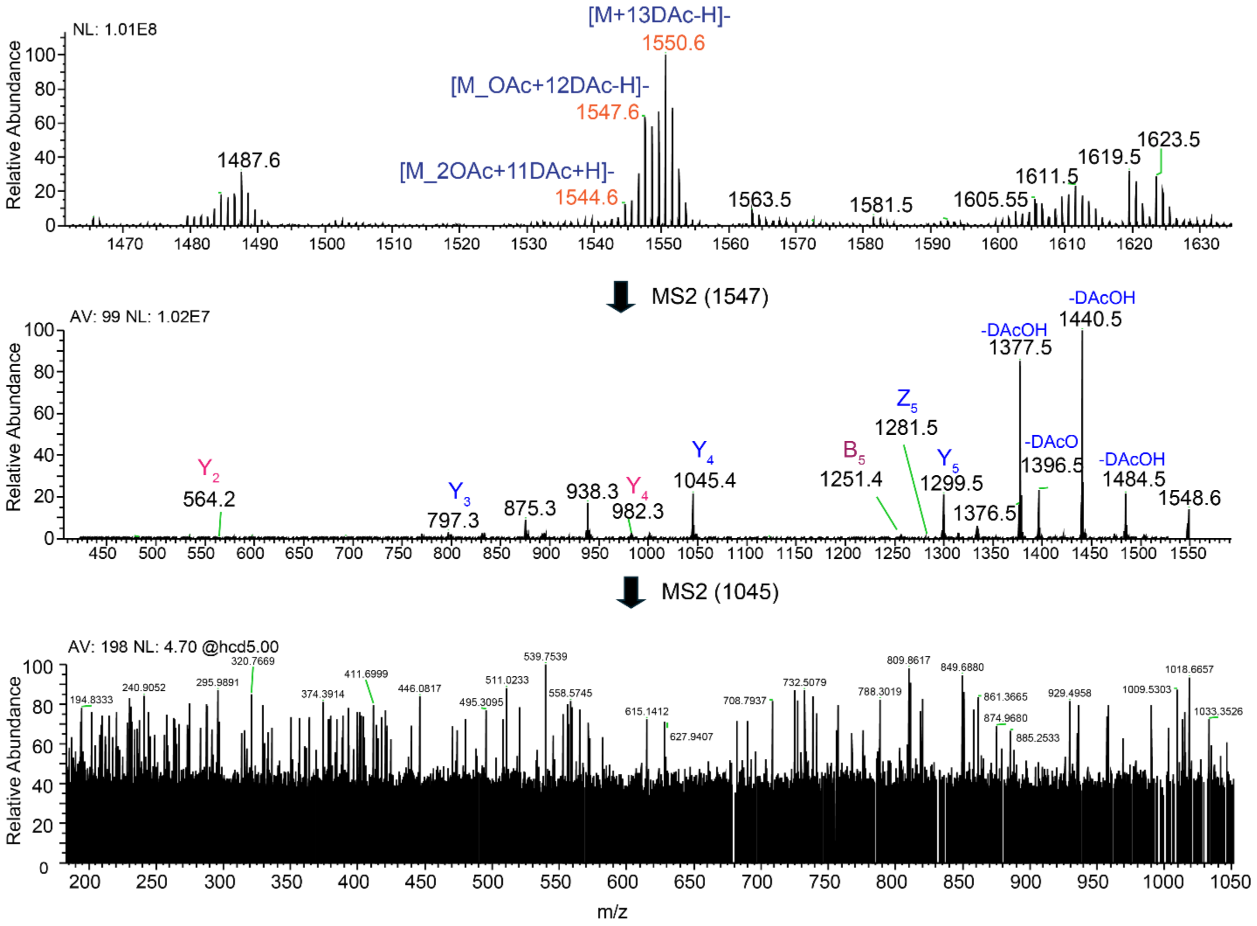


**Supplementary Figure 8:** ESI-MS$^n$ spectrum of perdeuteroacetylated, monoacetylated RG-I DP6 (top) in negative ion mode. The MS$^2$ spectrum of m/z 1547 [M-H]$^-$ parent ion (middle). The MS3 was obtained from the m/z= 1045 molecular ion (bottom). NL: normalized level, AV: averaged number of scans. Non-labeled peaks composition is not determined.

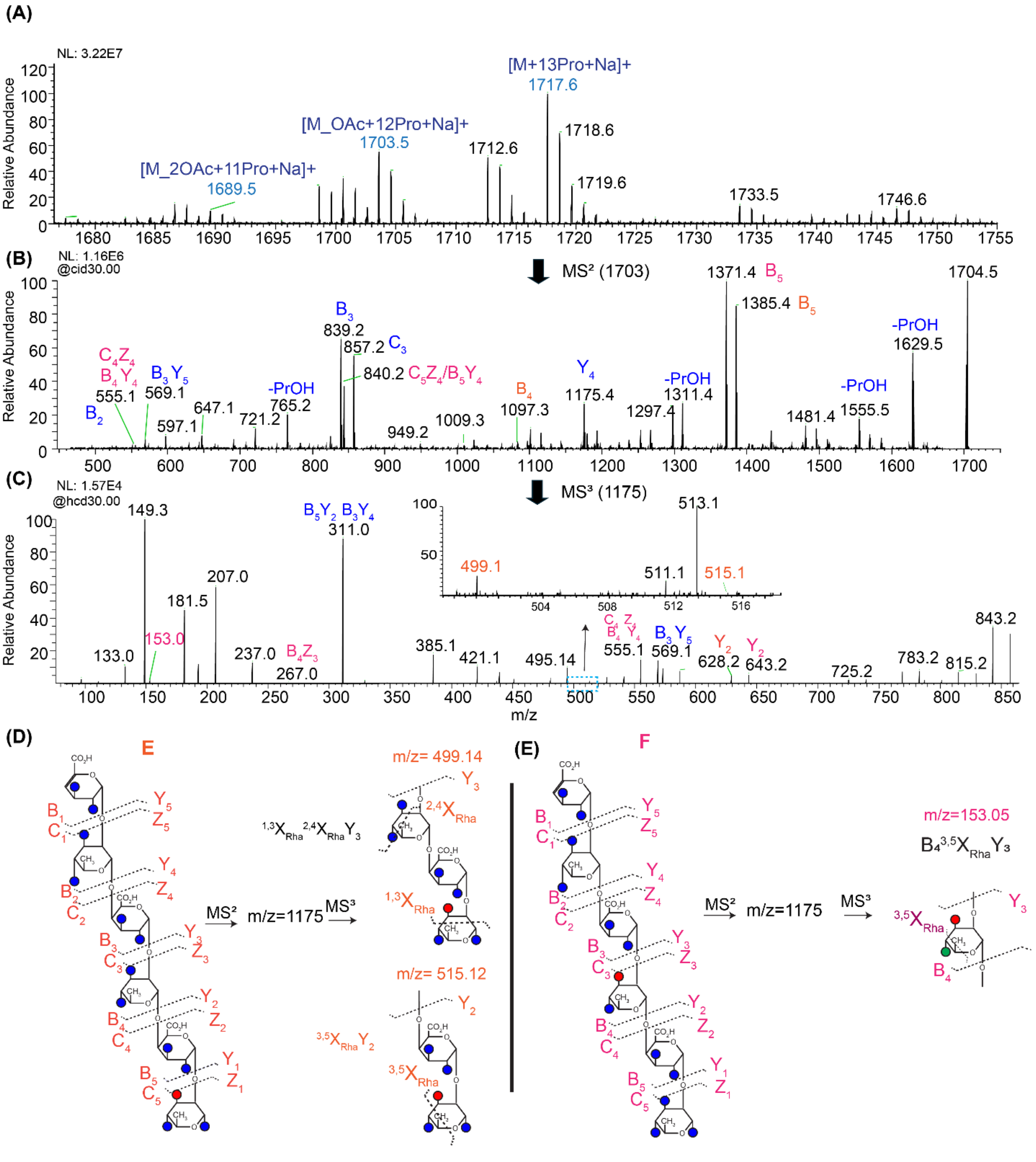


**Supplementary Figure 9**: **(A)** ESI mass spectra of the propionylated RG-I DP6 in positive ion mode. **(B)** $MS^2$ spectrum of the $[M\_OAc+12Pro+Na]^+$ ion *m/z*=1703, Pro denotes propionyl groups. **(C)** $MS^3$ spectrum of *m/z* = 1175. The inset shows the zoom in region mark in blue box. **(D)** Fragmentation pattern and the diagnostic cross ring fragment observed for structure E, where acetylation is on the 3-O position of the reducing end rhamnose. **(E)** Diagnostic cross-ring cleavage for structure F *m/z* = 153, where acetylation on the 3-O position of the second rhamnose. The blue circle on the structures denotes propionyl groups and red circle shows the acetyl position. The fragments correspond to Structure E labeled in orange, and for Structure F labeled in pink, common fragments for both E and F are labeled in blue. NL: Normally, the structures of the non-labeled peaks could not be determined.

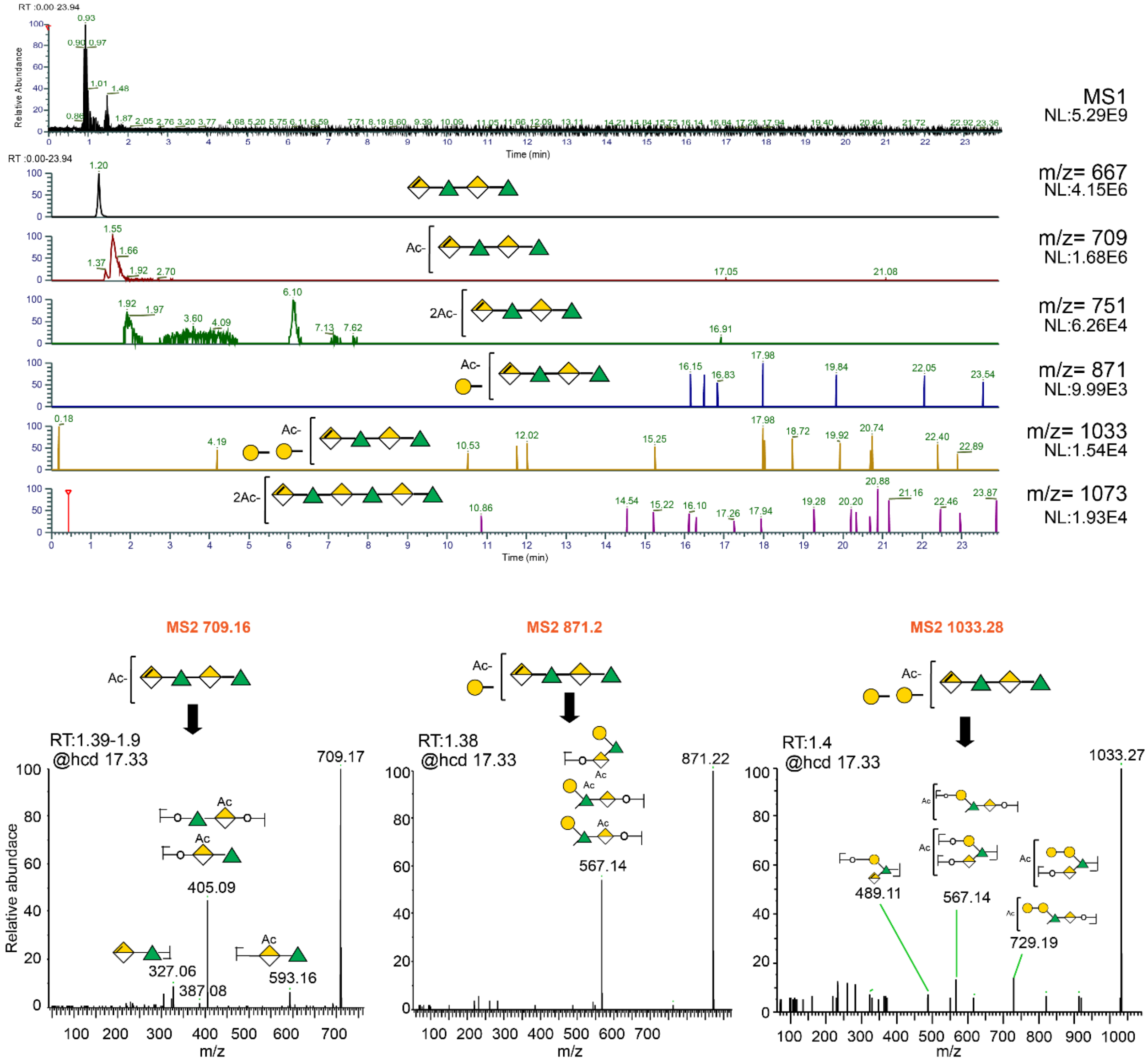


**Supplementary Figure 10:** LC elution profile and the LC-ESI-MS2 spectra of the native (underivatized) celery RG-I oligomers in positive ion mode [M+Na]+; The top panel shows LC elution profile of the m/z= 709 mono acetylated RG-I DP4 and m/z= 871 mono acetylated RG-I with one galactose branching.

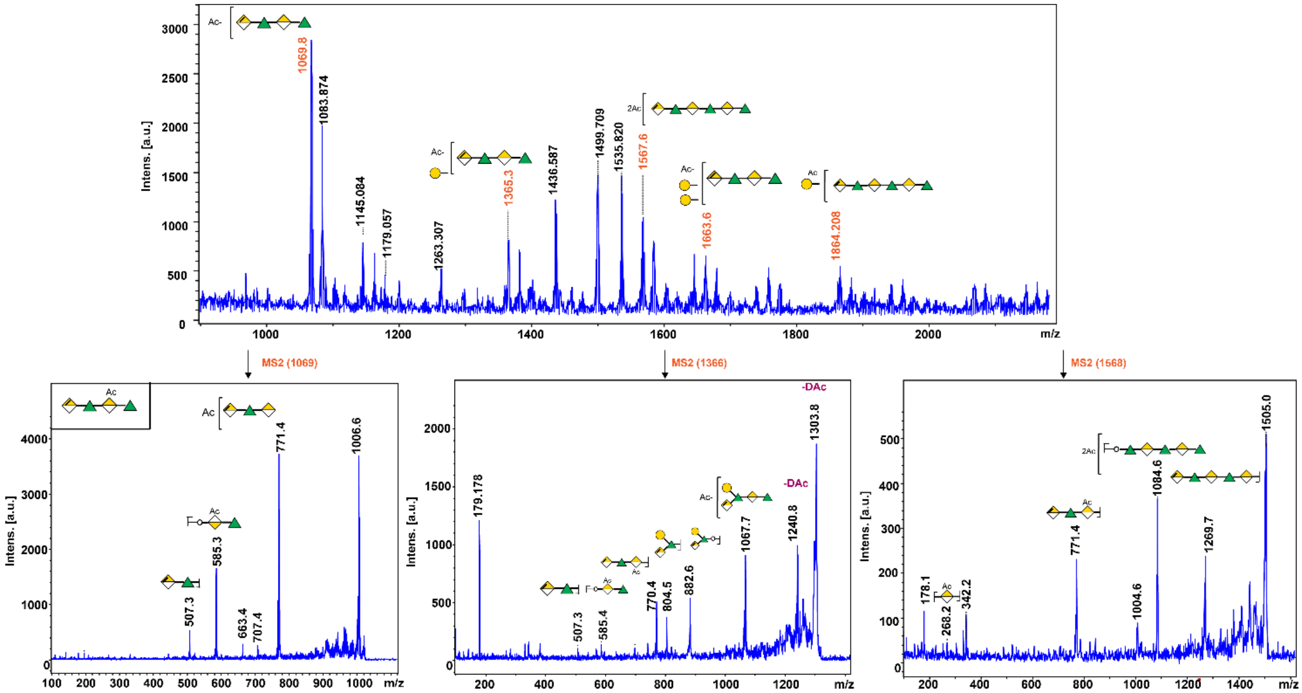


**Supplementary Figure 11:** MALDI-TOF MS spectra of deuteroacetylated oligomers derived from celery RG-I (top). The $MS^2$ spectra of different structures of RG-I; m/z =1069 mono acetylated RG-I DP4, m/z 1366 mono acetylated RG-I DP4 with galactose branching, m/z 1567 = diacetylated RG-I DP6 (bottom).

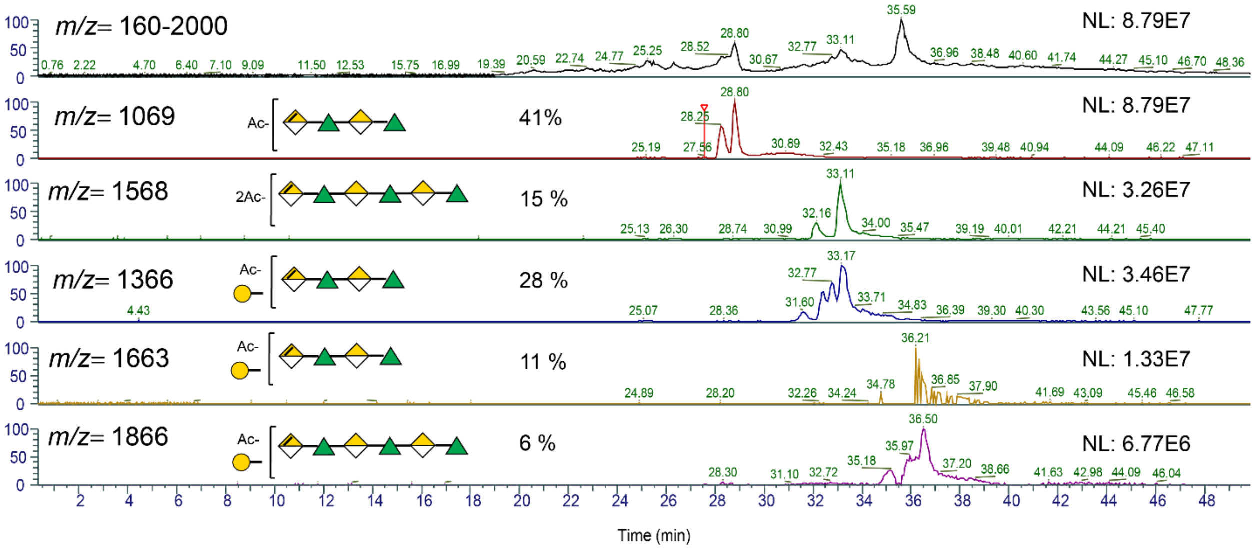


**Supplementary Figure 12**: Top: Full LC-MS acquisition of the mixture of deuteroacetylated oligomers derived from celery RG-I. Bottom: Extracted *m/z* values of different structural oligomers. Each structure shows different elution profile based on the degree of polymerization, degree of acetylation, branching pattern, and the length of the branching which related to the hydrophobicity of the oligomer. More hydrophobic the oligomer more time in the C18 reverse phase column and later elution. The ambiguous linkage positions of the acetyl group and the galactose reissue are shown in the brackets. The relative percentage of each structure shown in the spectra, calculated based on the peak intensity. NL: normalized intensity level.

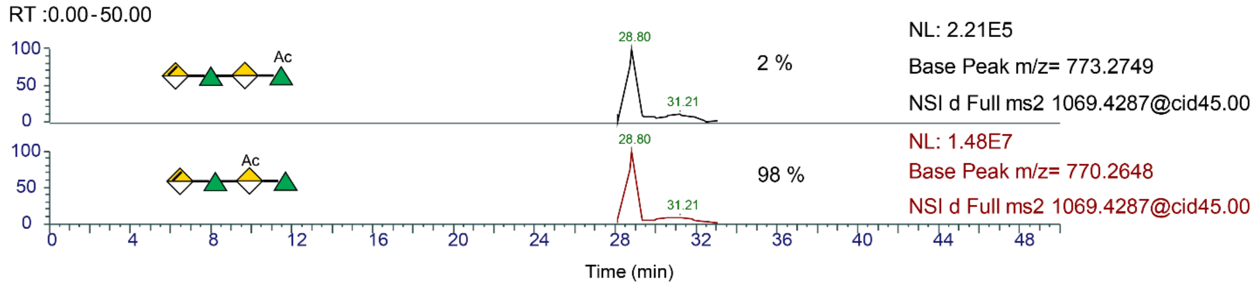


**Supplementary Figure 13:** LC Extraction of $B_3$ fragment; *m/z* =770 and *m/z* =773 derived from the MS2 of m/z= 1069 (RG-I_OAC DP4). The m/z=773 characteristic to the acetylation on the rhamnose residue and the *m/z* =770 corresponds to the acetylation in the second GalA residue. The relative percentage of each isomer are shown to be calculated based on the relative intensity of the peaks.

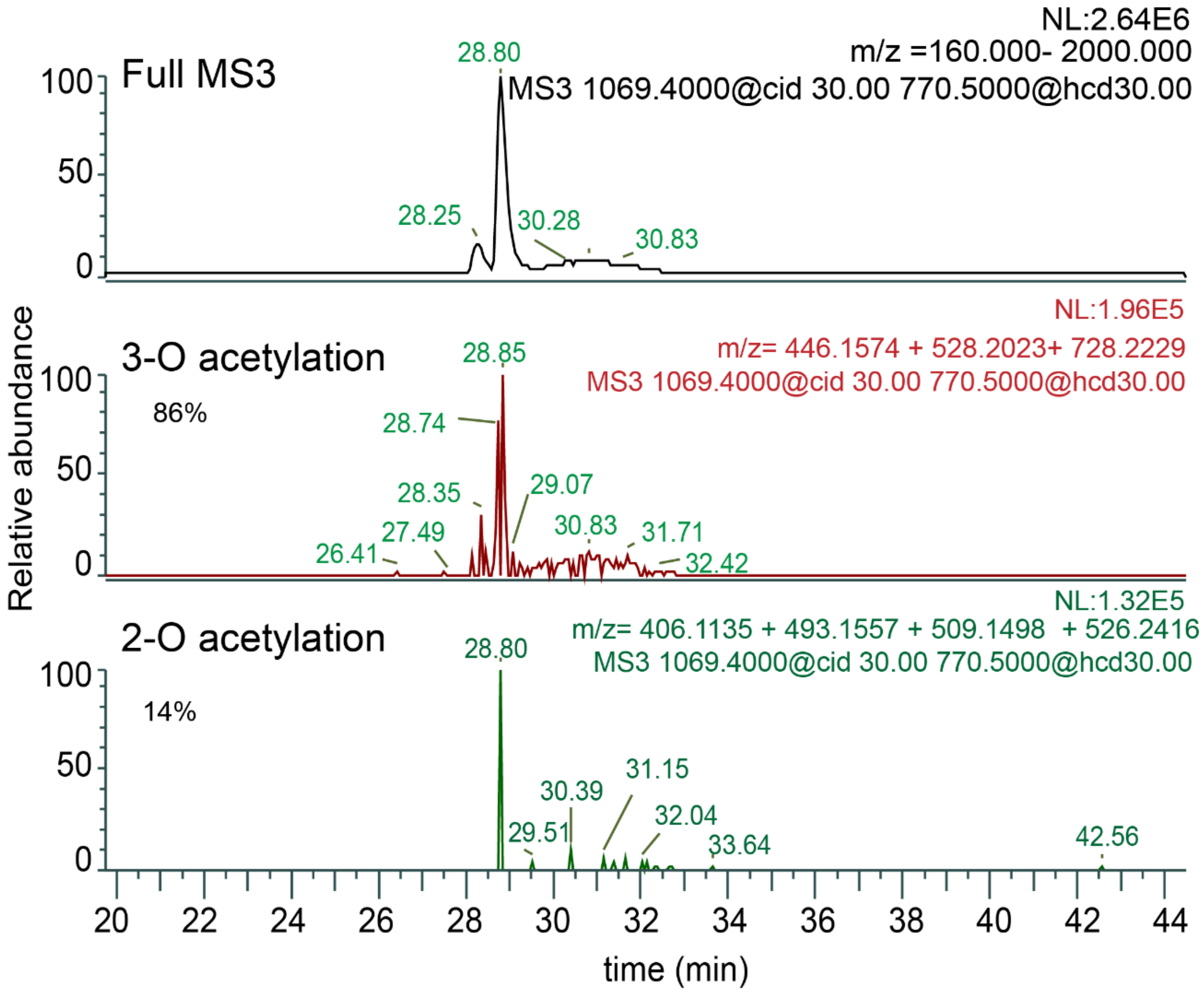


**Supplementary Figure 14:** Extracted *m/z* values for 3-OAc and 2-OAc from the full scan acquisition. The qualitative relative percentage of each structure shown in the spectra, calculated based on the peak intensities. Approximately the 3-OAc is 86% and 2-OAc is 14%. NL: normalized intensity level.

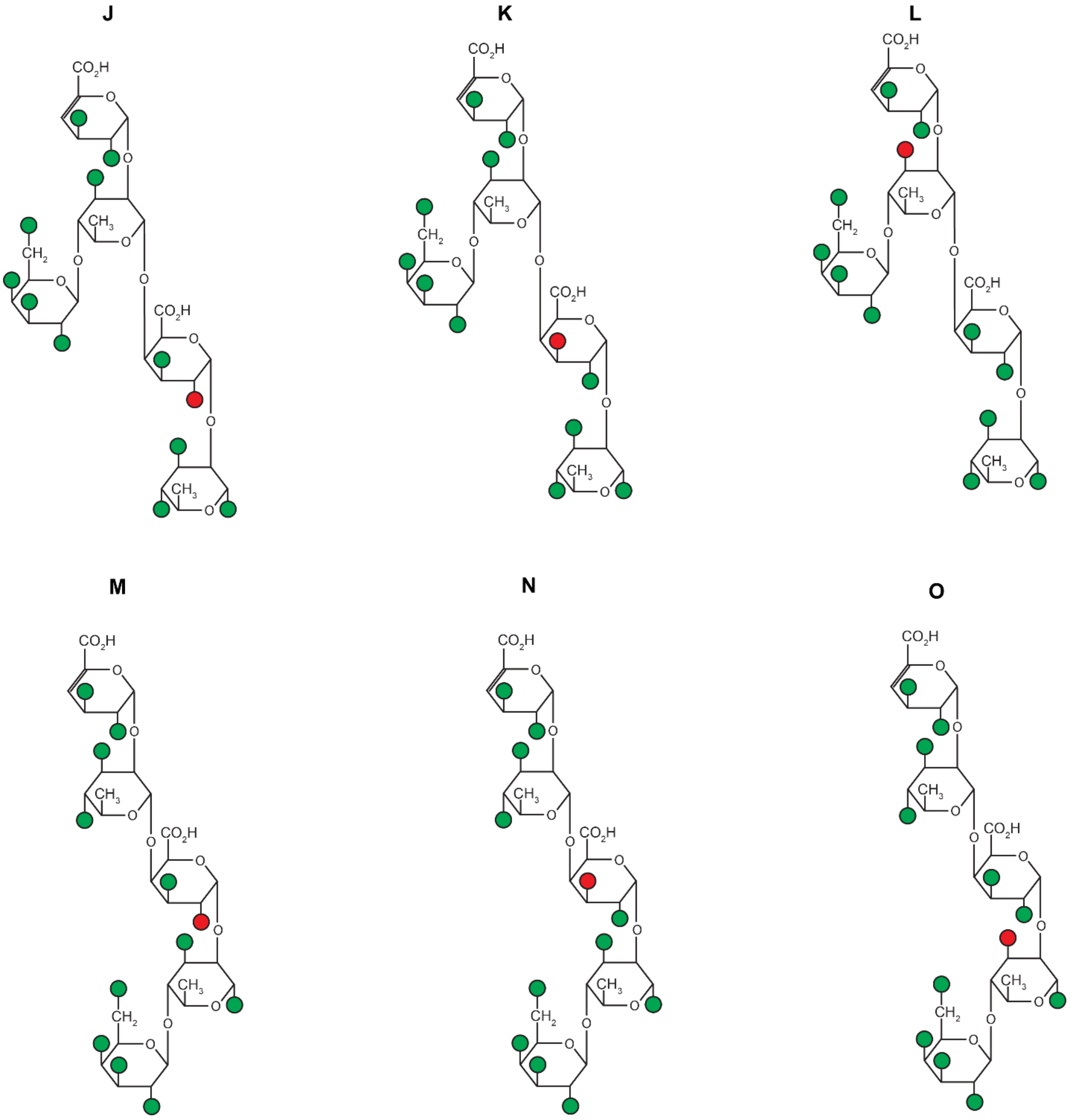


**Supplementary Figure 15:** The possible structures observed for the molecular ion *m/z*=1366; which comes from mono acetylated RG-I DP4 with one galactose branching. The structures are resulting from different acetylation position on the rhamnose and galacturonic acid, as well as the possible positions of the galactose residue. Based on the NMR the branching is on the rhamnose residue and deduce the possible structures,

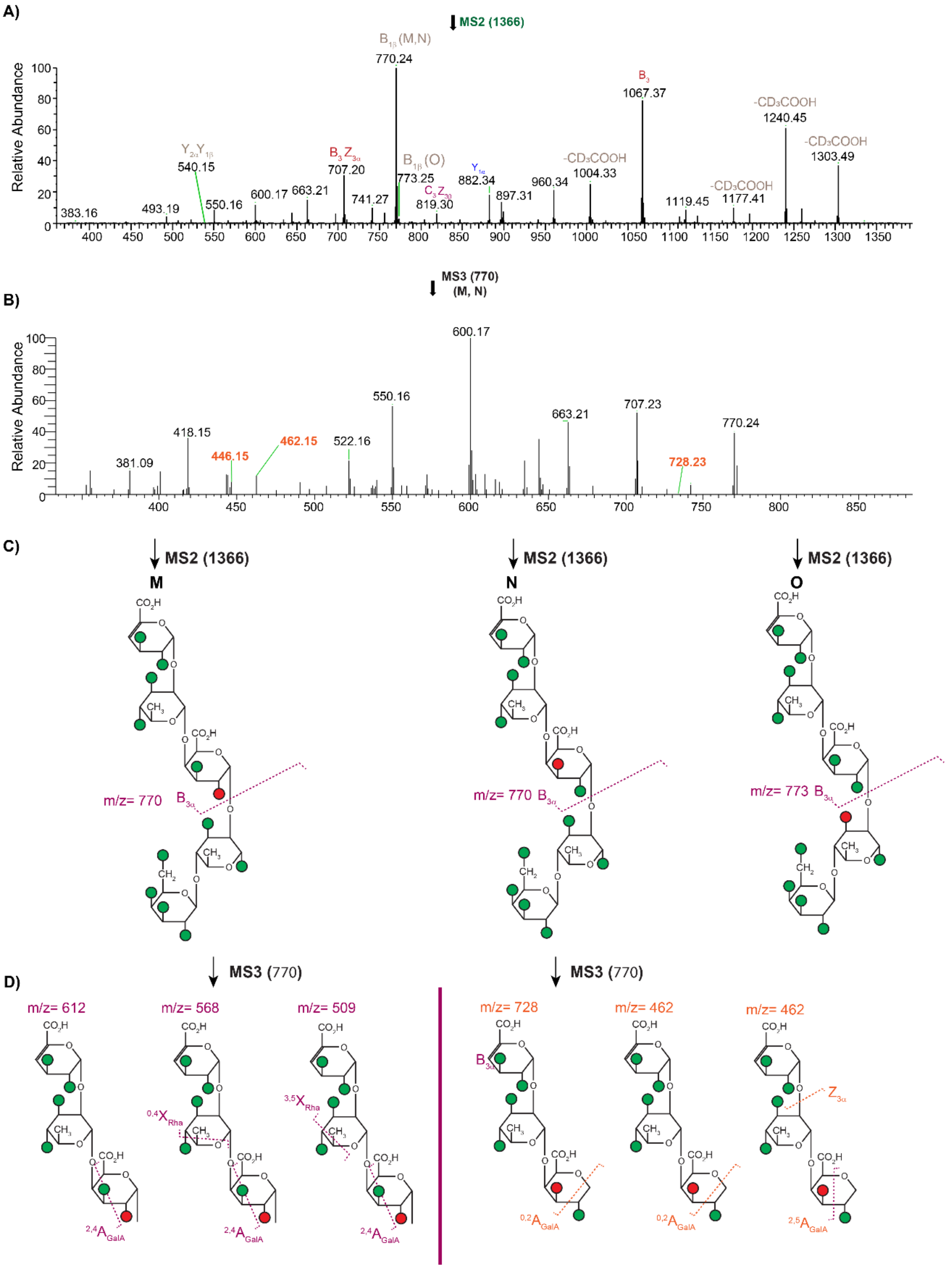


**Supplementary Figure 16:** **A)** $MS^2$ spectrum of *m/z*=1366 [M+Na]+ **B)** $MS^3$ spectrum of *m/z*= 770 (for M and N isomers) **C**) The M, N and O structures and cleavages observed in the MS spectra are labeled. The red circle denotes the acetyl groups, and the green circles are deuteroacetylated groups. **D)** Structure of the diagnostic cross-ring fragments observed in the $MS^3$ spectra for 2-O acetylation structure M and 3-O acetylation for structure N.

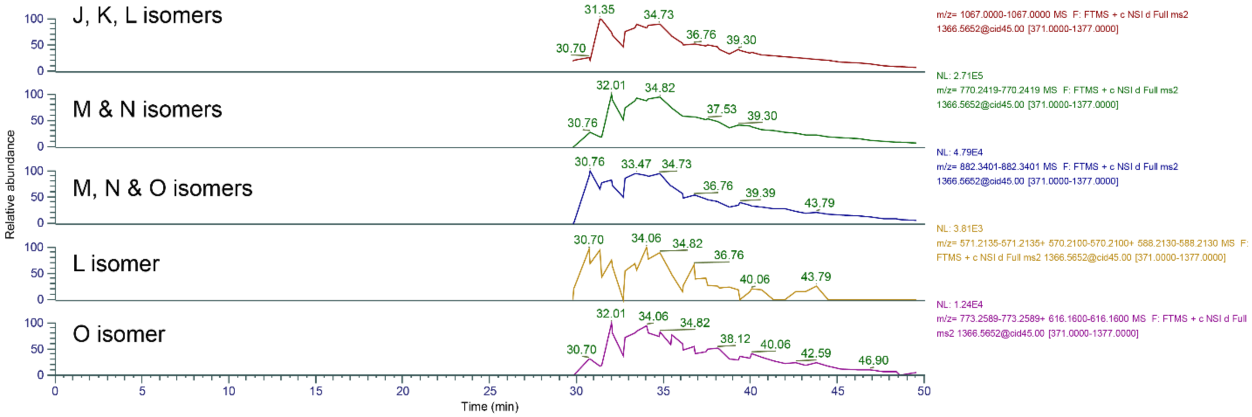


**Supplementary Figure 17:** Extraction of m/z values that are characteristic of different isomers from the full scan acquisition of $MS^2$ *m/z* =1366.

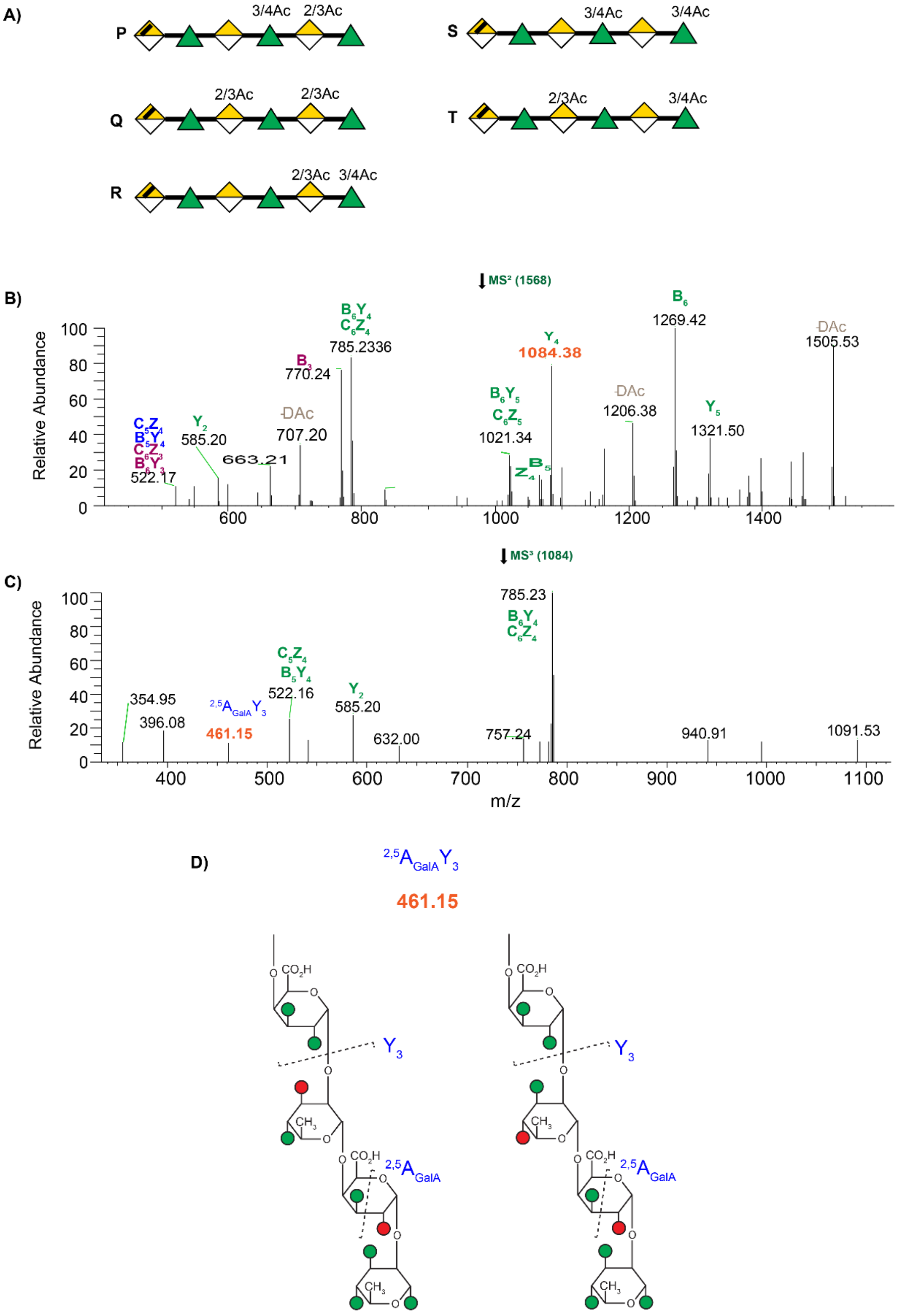


**Supplementary Figure 18: A)** Possible structures P, Q, R, S and T deduced from the MS2 spectra of the m/z= 1568 **B**) MS$^2$ spectrum of *m/z*=1568 [M+Na]+, perdeuteroacytalated linear DP6 RG-I_2OAc **C)** MS$^3$ spectrum of *m/z*= 1084) **D)** Diagnostic cross-ring cleavage observed in the MS spectra which pinpoints the GalA 3-O acetylation on the structure P. The red circle denotes the acetyl groups, and the green circles are deuteroacetylated groups

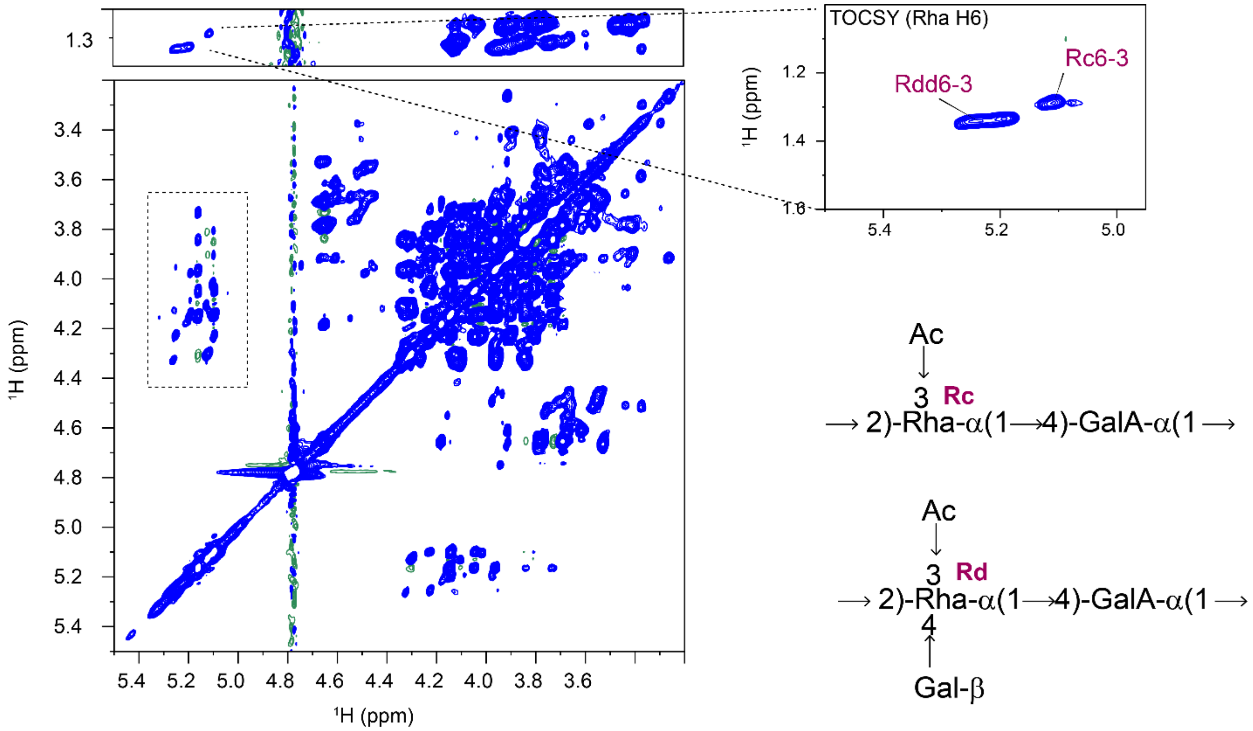


**Supplementary Figure 19:** $^1$H -$^1$H 2D TOCSY spectrum of native RG-I celery (undigested). TOCSY allows the observation of correlations between the anomeric proton (H1) and all other protons (H2–H6) within the rhamnose C6 methyl group (–$CH_3$) which appears at ~1.3 ppm. These chemical shifts Zoomed in shows the change in the position of the H6–H3 cross-peak compared to branched and non-branched acetylated rhamnose.

**Supplementary Table 1:** NMR Acquisition Parameters

| | Acquisition parameters at 25 °C at 800 MHz | | | | | | | | | | |
|---|---|---|---|---|---|---|---|---|---|---|---|
| Sample | Experiments | d1 (s) | NS | td2 | td1 | aq2 (s) | aq1 (s) | sw2 (ppm) | sw1 (ppm) | Tm (ms) | Exp Time |
| RG-I OAc DP6 | 1D $^1$H | 1.5/ 30s | 8 | 65536 | | 2.03 | - | 20.2 | - | - | 38s/10 min |
| | COSY | 1.5 | 4 | 2048 | 256 | 0.10 | 0.026 | 12 | 12 | - | 30min |
| | HSQC | 1.5 | 16 | 2048 | 256 | 0.10 | 0.003 | 12 | 160 | - | 2hr |
| Celery RG-I | 1D $^1$H | 1.0 | 4 | 65536 | - | 2.0 | - | 20.1 | - | - | 21s |
| | TOCSY | 1.5 | 16 | 2048 | 256 | 0.13 | 0.05 | 9.7 | 3 | 120 | 2hr |

**Supplementary Table 2:** $^{1}$H and $^{13}$C Chemical shifts (ppm) of RG-I samples.

RG-I_OAc DP6

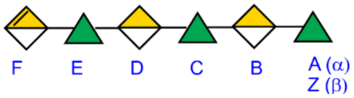


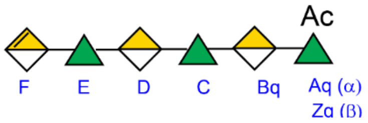


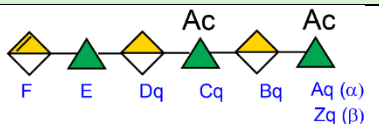


| Residue | 1 | 2 | 3 | 4 | 5 | 6 |
|---|---|---|---|---|---|---|
| E | 5.32<br>101.2 | 4.32<br>/ | / | / | / | / |
| C | 5.26<br>101.2 | 4.13<br>/ | / | / | / | / |
| Cq | 5.23<br>101.8 | 4.23<br>/ | / | / | / | / |
| A(α) | 5.22<br>94.32 | 3.98 | / | / | / | / |
| Z (β) | 4.93<br>96.6 | 3.99<br>/ | | | | |
| Aq(α) | 5.18<br>94.9 | / | | | | |
| Zq(β) | 5.00<br>96.2 | / | | | | |
| F | 5.11<br>100.5 | 3.81<br>/ | / | / | / | / |
| B | 5.05<br>100.4 | 3.93<br>/ | / | / | / | / |
| D | 5.01<br>100.1 | 3.94<br>/ | / | / | / | / |
| Bq | 4.97<br>101.9 | 3.91<br>/ | / | / | / | / |
| Dq | 4.96<br>101.7 | 3.92<br>/ | / | / | / | / |
| RG-I celery | | | | | | |
| 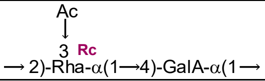 | / | / | 5.11 | / | / | 1.28 |
| 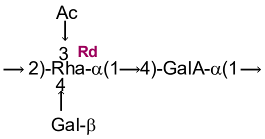 | / | / | 5.19 | / | / | 1.33 |

/, not determined

**Supplementary Table 3:** Diagnostic cross-ring fragments observed in the $MS^3$ (m/z 770) $[M+Na]^+$ spectra for 3-OAc and 2-OAc.

Diagnostic cross Ring in MS2 (1069)-> MS3 (770)

| Acetyl position | Observed m/z | Theoretical m/z | \|Δ mass difference\| | Cleavage |
|---|---|---|---|---|
| 3-OAc | 446.1674 | 446.1619 | 0.0055 | $^{2,5}A_{GalA}Z$ |
| | 528.2123 | 528.2156 | 0.00329 | $^{2,4}X_{GalA}$ |
| | 728.2329 | 728.2477 | 0.0148 | $^{0,2}A_{GalA}$ |
| | 409.2206 | 409.1858 | 0.0348 | $^{0,2}X_{GalA}$ |
| 2-OAc | 406.1235 | 406.1306 | 0.0071 | $^{0,2}X_{GalA}$ |
| | 493.1657 | 493.1706 | 0.0049 | $^{2,4}A_{GalA}{}^{0,3}X_{Rha}$ |
| | 509.1598 | 509.1655 | 0.0057 | $^{2,4}A_{GalA}{}^{3,5}X_{Rha}$ |
| | 526.2516 | 526.2364 | 0.0152 | $^{2,4}A_{GalA}{}^{0,3}X_{GalA}$ |
| 3Ac/2Ac | 462.1627 | 462.1568 | 0.0059 | $^{0,2}A_{GalA}Z$ |

**Supplementary Table 4:** Glycosidic cleavage fragments observed in the MS2 spectra of m/z 1366 characteristics for different isomers. The green shading shows the cleavage observed for each isomer. m/z=1067, 819 are only for J, K and L isomers. m/z= 770 is only for M and N isomer.

| MS2 (m/z= 1366) | | | | | | | | |
|---|---|---|---|---|---|---|---|---|
| Observed mass [M+Na]+ | Theoretical [M+Na]+ mass | Mass difference (Da) | J 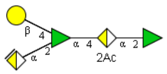 | K 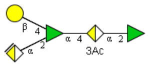 | L 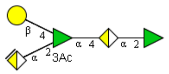 | M 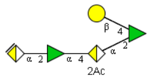 | N 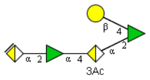 | O 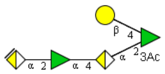 |
| 1067.3775 | 1067.39 | 0.0125 | $B_3$ | $B_3$ | $B_3$ | | | |
| 882.3401 | 882.3589 | 0.0188 | | | | $Y_{2\alpha}$ | $Y_{2\alpha}$ | $Y_{2\alpha}$ |
| 819.301 | 819.3190 | 0.018 | $B_3Y_{3\beta}/C_3Z_{3\beta}$ | $B_3Y_{3\beta}/C_3Z_{3\beta}$ | $B_3Y_{3\beta}/C_3Z_{3\beta}$ | | | |
| 770.2419 | 770.2583 | 0.0164 | | | | $B_{3\alpha}$ | $B_{3\alpha}$ | |
| 773.2589 | 773.2771 | 0.0182 | | | | | | $B_{3\alpha}$ |
| 707.2030 | 707.2183 | 0.0153 | $B_3Z_3$ | $B_3Z_3$ | $B_3Z_3$ | | | |
| 588.2133 | 588.2368 | 0.0235 | | | $Y_2$ | | | |
| 571.2134 | 571.2387 | 0.0253 | | | $C_2Y_{3\beta}$ | | | |
| 570.2119 | 570.2262 | 0.0143 | | | $Z_2$ | | | |
| 540.1553 | 540.1885 | 0.0332 | | | | $Y_{2\alpha}$ $Y_{1\beta}$ | $Y_{2\alpha}$ $Y_{1\beta}$ | $Y_{2\alpha}$ $Y_{1\beta}$ |
| 585.2047 | 585.2179 | 0.0132 | Y2 | Y2 | | | | |
| 556.2344 | 556.2469 | 0.0125 | $C_2Z_{3\beta}$ | $C_2Z_{3\beta}$ | | | | |
| 567.1946 | 567.2074 | 0.0128 | $Z_{16}$ | $Z_2$ | | | | |
| 574.3351 | 574.2575 | 0.0776 | $C_2Y_{1\beta}$ | $C_2Y_{1\beta}$ | | | | |

**Supplementary Table 5:** Glycosidic cleavages and cross-ring fragments observed in the $MS^3$ spectra of m/z= 1067 and m/z= 770. The m/z=1067 shows diagnostic fragments for J, K and L isomers. The m/z=770 shows diagnostic fragments correspond to the M and N isomers. (x) denotes common cross ring fragments observed in two or more isomers. The diagnostic cleavages only present in the L isomer shaded in green

| Observed mass [M+Na]+ | Theoretical mass [M+Na]+ | Difference (Da) | J 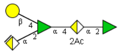  | K 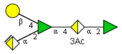  | L 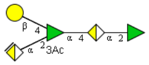  |
|---|---|---|---|---|---|
| MS3=1067 | | | | | |
| Glycosidic Cleavages | | | | | |
| 383.1607 | 383.1702 | 0.0095 | $C_{1\alpha}$ | $C_{1\alpha}$ | $C_{1\alpha}$ |
| 444.103 | 444.1463 | 0.0433 | $B_2Z_{3\alpha}$ | $B_2Z_{3\alpha}$ | |
| 480.127 | 480.1674 | 0.0404 | $C_2Y_{3\ \alpha}$ | $C_2Y_{3\ \alpha}$ | |
| 538.227 | 538.2364 | 0.0094 | $B_2Z_{3\beta}$ | $B_2\ Z_{3\beta}$ | |
| 553.256 | 553.2281 | 0.0279 | | | $C_2Z_{3\beta}$ /$B_2Y_{3\beta}$ |
| 556.234 | 556.2469 | 0.0129 | $C_2Z_{3\beta}$ | $C_2Z_{3\beta}$ | |
| 571.206 | 571.2387 | 0.0327 | | | $C_2Y_{3\beta}$ |
| 574.242 | 574.2575 | 0.0155 | $C_3Y_{3\beta}$ | $C_2Y_{3\beta}$ | |
| 804.308 | 804.3272 | 0.0192 | B2 | B2 | |
| 822.32 | 822.3378 | 0.0178 | C2 | C2 | |
| Cross-ring cleavages | | | | | |
| 355.1247 | 355.127 | 0.0023 | x | x | x |
| 357.1005 | 357.1063 | 0.0058 | x | x | x |
| 361.1968 | 361.1012 | 0.0956 | x | x | x |
| 363.1278 | 363.1169 | 0.0109 | x | x | |
| 364.1442 | 364.1201 | 0.0241 | x | | x |
| 365.1496 | 365.1596 | 0.01 | x | x | x |
| 376.1233 | 376.1201 | 0.0032 | x | x | |
| 380.1386 | 380.1150 | 0.0236 | x | | |
| 383.1607 | 383.1702 | 0.0095 | x | x | x |
| 391.1155 | 391.1118 | 0.0037 | x | x | x |
| 399.1086 | 399.1169 | 0.0083 | | x | x |
| 403.1318 | 403.1482 | 0.0164 | x | x | x |
| 416.1925 | 416.1514 | 0.0411 | x | x | |
| 419.0376 | 419.1067 | 0.0691 | x | x | x |
| 423.191 | 423.2015 | 0.0105 | $^{3,5}A_{Rha}$ | $^{3,5}A_{Rha}$ | $^{3,5}A_{Rha}$ |
| 491.112 | 491.1913 | 0.0793 | | | $B^{0,4}X_{Rha}Z$ |
| 495.211 | 495.2226 | 0.0116 | | | $^{2,5}A_{Rha}$ |
| 507.174 | 507.223 | 0.049 | | | $^{1,5}A_{Rha}Z$ |
| 668.248 | 668.2630 | 0.015 | | | $^{3,5}A_{GalA}Z$ |
| 717.296 | 717.2873 | 0.0087 | | | $B^{1,3}X_{Rha}$ |
| 745.255 | 745.2822 | 0.0272 | | | $^{1,4}A_{Rha}$ |
| 1012.36 | 1012.4126 | 0.0526 | x | | |

| Observed mass [M+Na]+ | Theoretical mass [M+Na]+ | Difference (Da) | M 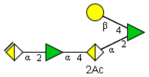  | N 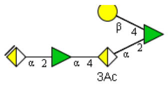  |
|---|---|---|---|---|
| MS3=770 | | | | |
| Cross-ring cleavages | | | | |
| 728.2363 | 728.2477 | 0.0114 | | $^{0,2}A_{GalA}$ |
| 462.1456 | 462.1568 | 0.0112 | | $^{0,2}A_{GalA}Z$ |
| 446.1509 | 4461619 | 0.011 | | $^{2,5}A_{GalA}Z$ |
| 607.2245 | 607.2023 | 0.0222 | | $^{2,5}A_{GalA}\ ^{1,3}X_{4uGalA}$ |
| 612.2025 | 612.2368 | 0.0343 | $^{2,4}A_{GalA}$ | |
| 610.2092 | 610.2211 | 0.0119 | $^{2,5}A_{GalA}\ ^{1,3}X_{4uGalA}$ | |
| 568.2353 | 568.2105 | 0.0248 | $^{2,4}A_{GalA}\ ^{0,4}X_{Rha}$ | |
| 509.1412 | 509.1655 | 0.0243 | $^{2,4}A_{GalA}\ ^{3,5}X_{Rha}$ | |
